# Awe is characterized as an ambivalent experience in the human behavior and cortex: integrated virtual reality-electroencephalogram study

**DOI:** 10.1101/2024.08.18.608520

**Authors:** Jinwoo Yi, Danny Dongyeop Han, Seung-Yeop Oh, Jiook Cha

## Abstract

Ambivalent feelings are a defining feature of awe, which has been understood as a possible source of its psychosocial benefits. However, due to the conventional unidimensional model of affective valence, behavior and neural representation of ambivalent feelings during awe remain elusive. To address this gap, we combined awe-inducing virtual reality clips, electroencephalogram, and a deep learning-based dimensionality reduction technique (*N* = 43). Behaviorally, awe ratings were precisely predicted by the duration and intensity of ambivalent feelings, not by single valence-related metrics. In the electrophysiological analysis, we identified latent neural space for each participant sharing valence representation structures across individuals and stimuli. In these spaces, ambivalent feelings during awe were distinctly represented from positive and negative ones, and the variability in their distinctiveness specifically predicted awe ratings. Additionally, frontal delta oscillations mainly engaged in differentiating valence representations. Our findings demonstrate that awe is fundamentally an ambivalent experience reflected in both behavior and electrophysiological activities. This work provides a new framework for understanding complex emotions and their neural underpinnings, with potential implications for affective neuroscience and relevant fields.

## Introduction

Awe, an emotion elicited by vast and enigmatic stimuli, both expands our minds and humbles us to our core. Understanding awe is crucial for unraveling the complexity of human emotional experiences and their impact on cognition, behavior, and well-being. Unlike simpler emotions, awe represents a unique interplay of cognitive expansion and self-diminishment, offering insights into how the human mind processes and integrates seemingly contradictory affective states. Unlike fear, awe is characterized by expanding one’s conceptual frameworks in an attempt to comprehend the mysterious. Psychologists define awe through two key dimensions: ‘perceived vastness’ and ‘a need for accommodation’ (Keltner & Haidt, 2003). For example, astronauts describe their awe during space flights as being overwhelmed by the vast scale of the universe compared to the smallness of Earth (i.e., perceived vastness) while also feeling a profound sense of beauty and fragility, leading to a realization of the urgent need to preserve this beauty (i.e., need for accommodation)(Yaden et al., 2016).

This multidimensionality of awe shapes its ambivalent nature, and this duality is expected to lead to significant psychiatric, psychosocial, and intellectual benefits, such as stress resilience, empathy, and non-egocentric perspectives (Jiang et al., 2024; Li et al., 2023; Monroy & Keltner, 2023; Rudd et al., 2012; Yin et al., 2024). Hence, as Keltner and Haidt (2003) argued, a comprehensive understanding of awe necessitates explaining its capacity to evoke both intensely positive and negative feelings.

However, despite its profound implications for human well-being, the ambivalent nature of awe remains underexplored in affective science. Current research struggles to capture awe’s inherent duality, often attempting to categorize it as simply positive or negative (Gordon et al., 2017; Guan et al., 2019; Piff et al., 2015; Takano & Nomura, 2022). This approach overlooks the key characteristic of awe — its capacity to evoke both positive and negative feelings simultaneously, prompting a fundamental reevaluation of our place in the world.

Two key methodological and theoretical bottlenecks hinder progress in understanding awe’s ambivalence. Firstly, traditional valence measurement tools, often relying on unidimensional scales (Russell, 2003), fail to capture the co-occurrence of positive and negative feelings. Both psychometric (An et al., 2017; Briesemeister et al., 2012) and neurobiological (Berridge, 2019; Lammel et al., 2012; Norman et al., 2011; Reynolds & Berridge, 2008) evidence suggest that a multidimensional model, representing positivity and negativity as distinct dimensions, offers a more accurate depiction of the valence system. For example, reward and aversion circuits, while both involving the ventral tegmental area (VTA), receive input from distinct brain regions: the laterodorsal tegmentum for reward and the lateral habenula for aversion (Lammel et al., 2012). This separation of functional neuroanatomy highlights the distinct pathways associated with positive and negative valence. Moreover, recent studies using negative awe-inducing images have demonstrated a higher co-occurrence of opposing valence than images evoking simple happiness or fear (Chaudhury et al., 2022). This finding suggests that traditional unidimensional valence scales may be insufficient for capturing the complex interplay of positive and negative feelings inherent in awe, underscoring the need for more nuanced measurement approaches.

Secondly, a lack of consensus on the neural representation of ambivalent feelings further complicates the study of awe. While the constructivist perspective posits that mixed feelings arise from rapid fluctuations between opposing valence systems (Barrett & Bliss-Moreau, 2009; Russell, 2017), emerging evidence suggests that ambivalent feelings may have distinct representations in the cortex (Man et al., 2017; Vaccaro et al., 2020). Human fMRI studies indicate that the posterior-anterior axis gradient within the right temporoparietal cortex is associated with valence co-occurrence during movie watching (Lettieri et al., 2019). The ventromedial prefrontal and anterior cingulate cortex also exhibit consistent neural patterns for ambivalent feelings (Vaccaro et al., 2024). Thus, it is required to assess how distinct neural representations mixed feelings during awe show for filling the knowledge gap regarding neural basis of ambivalent feelings.

To address these challenges, in this study, we combine the immersive power of 360° virtual reality (VR) with the high temporal resolution of electroencephalography (EEG) and robust deep learning-based dimensionality reduction analysis. VR offers unprecedented ecological validity crucial for awe research compared with conventional image and movie presentation on a screen (Chirico et al., 2016; Silvia et al., 2015), allowing participants to experience vast and enigmatic stimuli in a more profound and immersive manner than traditional methods (Chirico et al., 2017; Chirico et al., 2018; Kahn & Cargile, 2021; Quesnel & Riecke, 2018). EEG captures the fine-grained temporal dynamics of brain activity, essential for understanding the rapid interplay of affective processes. For instance, the insula cortex synthesizes bodily signals within a time window of about 125ms to create a global affective representation (Picard & Craig, 2009; Vaccaro et al., 2020; Wittmann, 2013), requiring a high temporal resolution to be measured. This approach enables precise tracking of the dynamic neural processes that underpin the complex affective states experienced during VR immersion.

To examine the distinct neural representation of ambivalent feelings during awe, we employed a contrastive learning-based deep learning approach to dimensionality reduction analysis – CEBRA (Schneider et al., 2023). It learns individual- and stimulus-specific latent neural spaces distinguishing EEG samples in terms of affective valence, efficiently exploring the representation of ambivalent feelings in addition to positive or negative ones. While dimensionality reduction offers a powerful way to reveal latent structures in neural data while preserving statistical power (Cunningham & Yu, 2014), traditional methods have struggled to uncover meaningful affect-related representations from non-invasive recordings. This difficulty stems from the highly nonlinear (Aftanas et al., 1998; Berridge, 2019; Viinikainen et al., 2010) and heterogeneous (Čeko et al., 2022; Lettieri et al., 2024) nature of human affect and low signal-to-noise ratio of non-invasive EEG recordings. By leveraging advanced algorithms based on nonlinear and supervised learning, our approach allows us to identify individual-specific latent neural spaces that capture both idiosyncratic valence representations and their commonalities across participants and stimuli. These individualized spaces, unlike conventional hand-crafted features such as frontal alpha asymmetry, provide a more nuanced and precise representation of the dynamic interplay of valence during awe. This integration of VR with EEG, analyzed through deep learning-based dimensionality reduction, forms a cohesive and innovative approach that bridges the gap between the subjective experience of awe and its neural substrates.

This study investigates the ambivalent nature of awe at both behavioral and neural levels, seeking to answer a fundamental question: How is awe, as a complex emotion characterized by the simultaneous experience of positive and negative feelings, represented in human behavior and the brain? To address this overarching question, we explore three specific inquiries: (1) Are awe ratings better predicted by ambivalence-related behavioral metrics than single valence measures? (2) Does ambivalent feeling during awe have a distinct neural representation, separable from positive and negative feelings? (3) Does the distinctiveness of ambivalent feeling representation in the neural space predict the intensity of awe ratings? By answering these questions, we aim to provide a more nuanced and comprehensive understanding of awe’s complex nature, revealing its fundamental connection to the interplay of positive and negative experiences.

## Results

### Overview of experimental protocols and behavioral data

Forty-three participants completed all experimental procedures (see **Fig 1a**). They viewed three awe-inducing clips (‘SP’ for ‘*Space*’, ‘CI’ for ‘*City*’, and ‘MO’ for ‘*Mountain*’) and one control clip (‘PA’ for ‘*Park*’) in a pseudo-random order, each lasting 120 seconds with sound. These clips were designed to evoke diverse awe experiences based on distinct awe-related factors and sensory features (see ‘Methods’ and **Supplementary Fig 1**). During watching movies, participants’ EEG signals were recorded, and they reported their valence dynamics through real-time keypress, indicating neutral, positive, negative, and ambivalent feelings (see **Fig 1b**). They held the key down for the duration of each affective feelings, and these ratings were synchronized with EEG samples. This approach allowed for a direct association between real-time affective response and neurophysiological data. After each video, they rated overall valence, arousal, awe, and motion sickness. Keypress valence ratings were used to calculate the duration of four valence feelings, while the post-trial valence ratings were applied to quantify the intensity of positive, ambivalent, and negative feelings.

**Fig 1.**
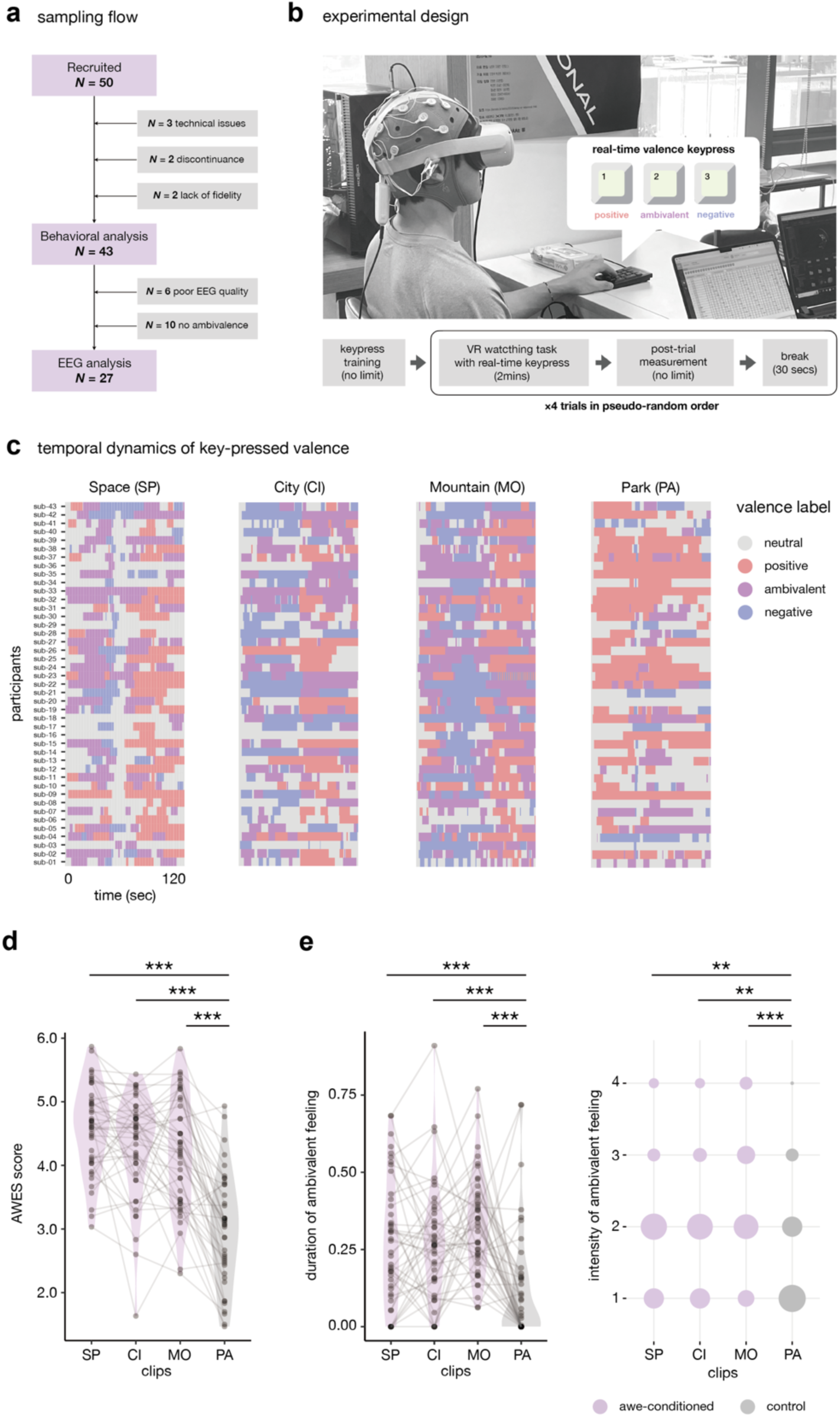
Valence dynamics, awe, and ambivalent feelings for VR clips. **a,** The sampling procedures, summarizing participant recruitment and selection for behavioral and EEG analysis. **b,** The integrated VR-EEG-real time valence rating protocol, showing the sequence of experimental activities. **c,** Valence dynamics rated by real-time keypress from 43 participants during exposure to each VR clip. **d,** AWES scores evaluated post-trial for each clip, reflecting the intensity of awe experiences. **e**, The duration (left) and intensity (right) of ambivalent feelings for each clip, comparing responses between awe-conditioned and control clips. \**P*_FDR_ < .05; \*\**P*_FDR_ < .01; \*\*\**P*_FDR_ < .001.

### Awe experiences are consistently associated with longer and stronger ambivalent feelings, but not with single-valence ones

We first explored participants’ valence dynamics ratings. Valence dynamics showed idiosyncratic temporal patterns (see **Fig 1c**), but they were also significantly intertwined to visual and acoustic features known to predict perceivers’ affective responses - e.g., color hue (Dael et al., 2016; Suk & Irtel, 2010), brightness (Kurt et al., 2017), and loudness (Thao et al., 2019)(see **Supplementary Table 1**). These results validate our real-time valence rating paradigm, showing that individual responses are systematically connected to affect-related sensory inputs, despite their idiosyncrasy.

Next, we examined if awe-inducing clips were associated with higher awe and ambivalence-related metrics. Participants reported significantly higher awe ratings for all awe clips than the control (SP vs. PA: Cohen’s *d* = 1.837, *P*_FDR_ = 1×10^-14^; CI vs. PA: Cohen’s *d* = 1.493, *P*_FDR_ = 3×10^-12^; MO vs. PA; Cohen’s *d* = 1.373, *P*_FDR_ = 2×10^-11^; see **Fig 1d**). The three awe clips also led to significantly longer and stronger ambivalent responses than the control (SP vs. PA: Cohen’s *d*_duration_ = .562, *P*_duration/FDR_ = .001, Cohen’s *d*_intensity_ = .451, *P*_intensity/FDR_ = .005; CI vs. PA: Cohen’s *d*_duration_ = .514, *P*_duration/FDR_ = .002, Cohen’s *d*_intensity_ = .449, *P*_intensity/FDR_ = .005; MO vs. PA: Cohen’s *d*_duration_ = .790, *P*_duration/FDR_ = 2×10^-5^, Cohen’s *d*_intensity_ = .780, *P*_intensity/FDR_ = 2×10^-5^; see **Fig 1e**). Contrarily, behavioral metrics of positive and negative feelings showed no consistently significant differences between conditions, except for the duration of positive feelings and arousal (see **Supplementary Table 2**). These results show the potential of awe experiences to evoke salient and persistent ambivalent feelings compared to simply positive or negative states.

### Ambivalent feelings predict the awe ratings more precisely than single valence metrics

We tested whether ambivalence-related metrics had greater predictive power for awe ratings than other single-valence features using 10 ratings from each clip and five baseline variables (sex, age, baseline positive and negative mood, and dispositional awe). The univariate linear mixed effects model analysis revealed that only the duration and intensity of ambivalent feelings and the intensity of positive feelings showed significant fixed effects (duration of ambivalent feelings: 𝛽 = .565, 95% CI = [.033, 1.097], *P* = .039; intensity of ambivalent feelings: 𝛽 = .220, 95% CI = [.090, .351], *P* = .001; intensity of positive feelings: 𝛽 = .094, 95% CI = [.007, .180], *P* = .035; see **Fig 2a**), confirming the unique contribution of emotional complexity to awe experiences.

**Fig 2.**
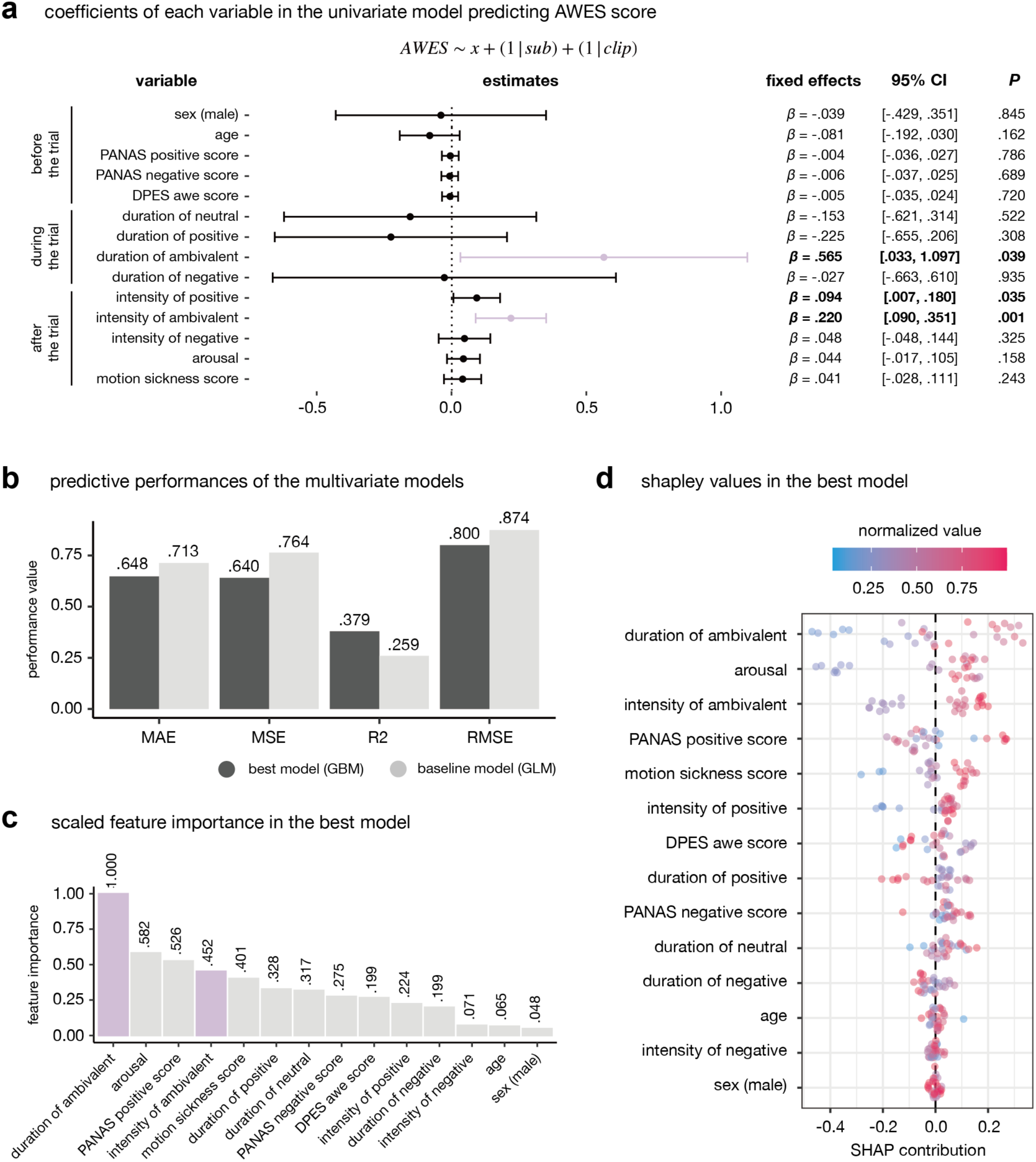
Association between behavioral metrics and awe ratings. **a,** Coefficients of each behavioral metric in the univariate linear mixed effect models. Error bars denote 95% confidence intervals of fixed effects. Purple bars show the estimates of ambivalence-related features. Bolded statistics indicate statistically significant results at *P* < .05. **b,** Comparison of predictive performances between AutoML-based best model (GBM) and linear regression model (GLM). **c,** Feature importance (scale from 0 to 1) of all behavioral features calculated from the best model. Purple bars show the importance of ambivalence-related features. **d,** Shapley values of all features computed from the best model.

To further explore the relationship between these variables, we employed an AutoML pipeline as multivariate analysis allowing for potential nonlinear interactions. Our best model, selected via 5-fold cross-validation, showed superior predictive performance than a linear regression model (see **Fig 2b**). In this model, ambivalent feelings’ duration and intensity ranked higher in feature importance than other valence metrics (see **Fig 2c**). Shapley values indicated a positive association between ambivalence and awe ratings (see **Fig 2d**). The results from both univariate and multivariate analyses provide compelling evidence that the ambivalent feeling is significantly related to stronger awe experiences.

### Overview of electrophysiological analysis

To examine neural representation of ambivalent feelings during awe and its relationship with awe ratings, we analyzed EEG signals and real-time valence keypress. In this analysis, we used data from 27 participants who showed EEG signals with acceptable qualities and responded ambivalent feelings in their keypress for all awe-inducing clips (see **Fig 1a**). These selection criteria were based on the premise that a consistent reporting of ambivalence across stimuli would provide a clearer signal for assessing its neural correlates. Given our interest of ambivalent feelings’ latent neural representation during awe and behavioral findings that the duration of ambivalent feelings was the most salient predictor of awe ratings, such sampling was required and expected not to introduce a sampling bias. Included and excluded participants did not differ significantly in key demographic measures or behavioral awe ratings (all *P*s > .05), suggesting that this selection did not introduce systematic bias (see **Supplementary Table 3**). We applied Short-time Fourier Transformation (STFT) to calculate band power across delta (1-4Hz), theta (4-8Hz), alpha (8-14Hz), beta (14-31Hz), and gamma (31-49Hz) frequency. Afterward, we performed dimensionality reduction to extract latent neural spaces from STFT features, using CEBRA (Schneider et al., 2023), a contrastive learning-based approach. CEBRA can capture complex, non-linear relationships in high-dimensional data while emphasizing distinctions between affective states, allowing for more effective isolation of neural representations specific to each state, including ambivalence. Indeed, CEBRA attracted EEG samples with the same valence state while repelling those with different valence states, revealing cohesive clusters for each valence state in the latent neural space (see **Fig 3a**). We independently trained CEBRA on STFT features of each participant-clip datum to identify individualized and stimulus-specific latent neural spaces.

**Fig 3.**
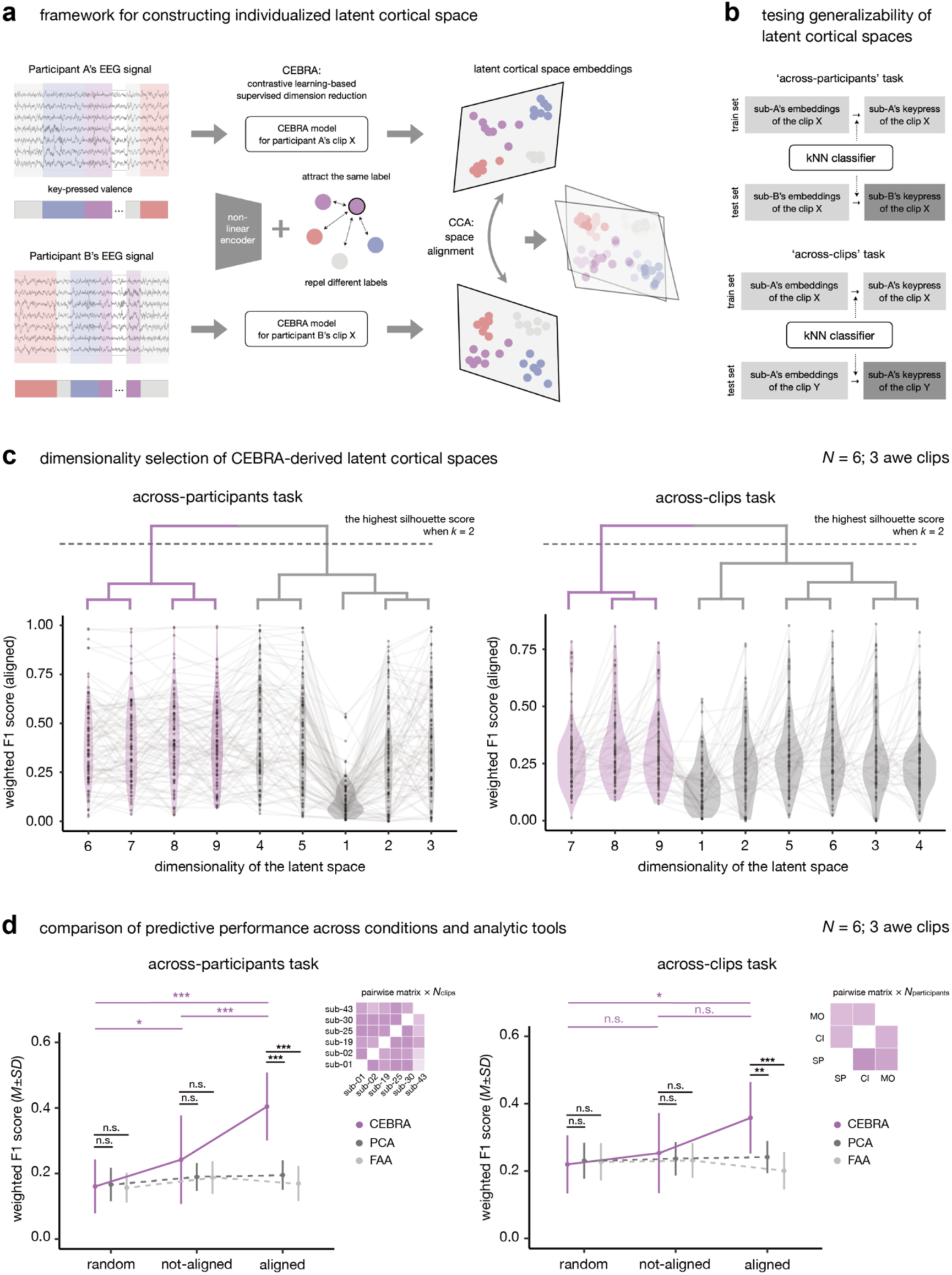
Identification of individualized latent neural spaces and evaluation of their generalizability. **a,** Schematic illustrating the extraction of individualized latent neural spaces using CEBRA and neural alignment. **b,** Schematic depicting the assessment of generalizability through pairwise predictive tasks across participants and sensory stimuli. **c,** Dimensionality selection for CEBRA-driven latent neural spaces. Hierarchical clustering analysis identified clusters with high performance in predicting individual differences (purple violin plots). **d,** Predictive performance of latent spaces derived from CEBRA, PCA, and FAA across three conditions. CEBRA-based embeddings consistently outperform PCA and FAA-based embeddings in predicting individual differences (black asterisks). Heatmaps visualize conceptual scheme of results from pairwise predictive tasks. Within CEBRA-based predictions, performance varies significantly across conditions (purple asterisks). Results are based on *N* = 6 participants reporting all valence types in all awe-inducing clips. \**P*_FDR_ < .05; \*\**P*_FDR_ < .01; \*\*\**P*_FDR_ < .001.

To assess if these individualized spaces share a common valence representation structure across participants and stimuli, we conducted two pairwise decoding tasks which predicted valence dynamics of other participants or other clips. Next, we tested if ambivalent feelings formed distinct neural clusters and if distinctiveness of these representations could predict awe ratings. Finally, we analyzed which EEG features mainly engaged in distinguishing valence types in the latent neural spaces.

### Aligned latent neural spaces share valence representation architecture across individuals and stimuli

To evaluate the generalizability and optimal dimensionality of individualized latent neural space constructed with CEBRA, we performed pairwise decoding analysis with two tasks (‘across-participants’ and ‘across-clips’) and three conditions (‘random’, ‘not-aligned’, and ‘aligned’). For the ‘across-participants’ task, a classifier was trained on participant A’s latent space embeddings to decode his valence dynamics and was tested on participant B’s embeddings for the same clip. In the ‘across-clips’ task, a classifier trained on latent space embeddings from clip X was tested on clip Y for the same participant (see **Fig 3b**). To facilitate multi-class classification, only six participants’ data for three awe-inducing clips were used, which included all valence categories in the keypress for all awe-inducing clips (see ‘Methods’ for detailed inclusion criteria). We assessed decoding performance under three conditions: classifiers trained on shuffled valence dynamics labels (‘random’), on personalized latent space embeddings without alignment (‘not-aligned’), and on latent embeddings aligned with test embeddings through canonical correlation analysis (‘aligned’). Alignment helps identify commonalities in individual neural spaces and enhances the interpretability of shared neural representations (Gallego et al., 2020; Safaie et al., 2023). Assuming that the optimal latent space is consistently dimensional across individuals and clips, achieving comparable predictive performance with the simplest dimensionality (see ‘Methods’ for rationales), we selected a 7D space for the optimal CEBRA-driven latent neural spaces (see **Fig 3c**). For a comparative baseline model, we performed the same analyses with principal component analysis (PCA)-driven latent spaces, and a 6D space was chosen for the optimal PCA-based embedding space (see **Supplementary Fig 2**).

We then compared decoding performances of the latent neural space across participants and sensory stimuli among CEBRA, PCA, and frontal alpha asymmetry (FAA), a traditional index of affective valence (see **Fig 3d**). In the ‘across-participants’ task, CEBRA-based classifiers performed significantly better than random chance (aligned vs. random: Cohen’s *d* = 1.122, *P*_FDR_ = 5×10^-17^; not-aligned vs. random: Cohen’s *d* = .249, *P*_FDR_ = .020). Aligned embeddings outperformed not-aligned ones (aligned vs. not-aligned: Cohen’s *d* = .548, *P*_FDR_ = 2×10^-6^). Also, they provided more accurate predictions than PCA-based (Cohen’s *d* = 1.093, *P*_FDR_ = 2×10^-16^) and FAA-based latent space embeddings (Cohen’s *d* = 1.115, *P*_FDR_ = 6×10^-17^) in aligned conditions, but not in random and not-aligned conditions (see Supplementary Table 4).

In the ‘across-clips’ task, only aligned CEBRA embeddings showed significant predictive power for valence dynamics across clips (aligned vs. random: Cohen’s *d* = .501, *P*_FDR_ = .015; not-aligned vs. random: Cohen’s *d* = .103, *P*_FDR_ = .540). Aligned embeddings outperformed not-aligned ones, although its significance did not reach a threshold (aligned vs. not-aligned: Cohen’s *d* = .283, *P*_FDR_ = .147). Aligned CEBRA embeddings more accurately predicted valence dynamics than PCA-based (Cohen’s *d* = .553, *P*_FDR_ = .006) and FAA-based embeddings (Cohen’s *d* = .807, *P*_FDR_ = 8×10^-5^), but not in random and not-aligned conditions (see **Supplementary Table 4**).

These findings suggest that aligned CEBRA-based latent neural spaces capture a shared architecture of valence representations, including ambivalent feelings, across individuals and stimuli, which could not be fully identified by conventional approaches such as PCA or FAA.

### The more distinctively ambivalent feelings are represented in the cortices, the more saliently individuals report awe

Using not-aligned latent neural spaces, we explored whether ambivalent feelings have distinct neural representations separable from representations of other single valence states. To quantify the cohesiveness of neural representations for each valence state, we calculated the silhouette coefficient of each valence-specific cluster in the latent neural spaces. A higher silhouette coefficient indicates that the corresponding valence type is represented as a more cohesive cluster in the latent space. Since the silhouette coefficient can be overestimated in supervised latent spaces as CEBRA-driven ones, we evaluated its statistical significance using 1,000 times permutation test (see **Fig 4a**).

**Fig 4.**
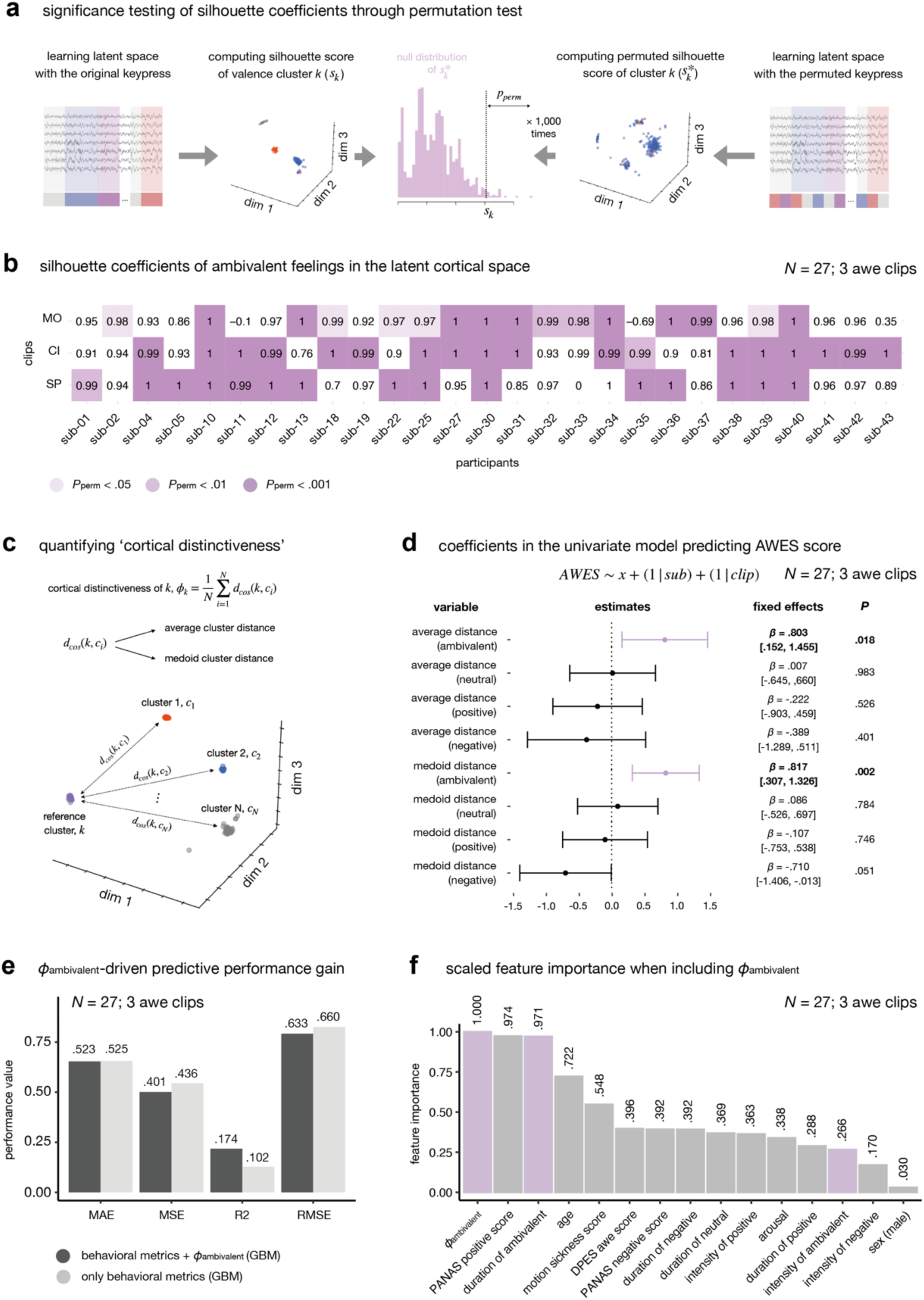
Individual variability of ambivalence-related latent neural representation and its predictive power on awe ratings. **a,** Schematic illustrating the assessment of statistical significance for silhouette coefficients through a permutation test. **b,** Silhouette coefficients of ambivalent valence clusters within individual latent neural spaces. Color intensity indicates the statistical significance of the silhouette coefficients. **c,** Schematic depicting the calculation of “cortical distinctiveness” for each valence cluster. See Equation (2) and (3) in ‘Methods’ section for annotations. **d,** Explanatory power of each valence cluster’s cortical distinctiveness in the univariate linear mixed-effects models. Error bar denotes 95% confidence interval of fixed effects. Purple bars show the estimates of ambivalence-related features. Bolded statistics indicate statistically significant results at *P* < .05. Notably, the cortical distinctiveness of ambivalent states (purple bars) significantly predicts awe ratings. **e,** Performance gain observed when cortical distinctiveness of ambivalent feelings was added to the behavioral prediction model. **f,** Scaled (0-1) feature importance of cortical distinctiveness of ambivalent feelings and behavioral features. In both, the cortical distinctiveness of ambivalent states (purple bars) emerges as a prominent predictor of awe. Results are based on *N* = 27 participants included in the electrophysiological analysis. \**P* < .05.

We observed that the silhouette coefficients of ambivalent clusters varied widely across participants (see **Fig 4b**), while single valence states showed relatively consistent significant silhouette coefficients (see **Supplementary** Fig 3). This individual variability in the ambivalence-related representations suggests that the integration of positive and negative experiences during awe may manifest differently across individuals. Concerned about potential confounding effects of cluster size on silhouette coefficients significance, we tested the correlation between the number of ambivalence-labeled EEG samples in the cluster (i.e., duration of ambivalent feelings) and the *P* values of silhouette coefficients for each clip. No significant correlations were found (SP: Pearson’s *R* = .221, 95% CI = [-.174, .554], Pearson’s *P* = .268; CI: Pearson’s *R* = .198, 95% CI = [-.197, .538], *P* = .323; MO: Pearson’s *R* = .126, 95% CI = [-.267, .483], *P* = .531), suggesting that the observed variability in silhouette coefficients for ambivalence is not merely due to differences in the duration of reported ambivalence.

To directly test the relationship between ambivalence-related neural representations and awe experiences, we then examined how spatial properties of valence clusters in the latent neural space relate to awe ratings. We developed a new metric called ‘cortical distinctiveness’ for cluster *k*, 𝜙*_k_*. It quantifies how distinguishable representations cluster *k* displays from other valence clusters by averaging cosine distance with other clusters (see **Fig 4c**). Higher 𝜙*_k_* indicates that the neural representation of the corresponding valence cluster *k* is more spatially separated from other valence representations within the individual’s latent neural space. Using this metric, we constructed four linear mixed-effect models including 𝜙 value of each valence cluster and two random intercepts of participants and clips. In these models, only 𝜙*_ambivalent_* significantly predicted awe intensity (𝛽 = .817, 95% CI = [.307, 1.326], *P* = .003; see **Fig 4d**). The predictive power of 𝜙*_ambivalent_* persisted even when using medoid cluster distance for calculating this metric (𝛽 = .803, 95% CI = [.152, 1.455], *P* = .018; see **Fig 4d**), demonstrating the robustness of the findings to the specific method of calculating cortical distinctiveness. As a control analysis, we conducted the same analysis with FAA values and found that FAA did not predict awe ratings significantly (𝛽 = .000, 95% CI = [-.004, .003], *P* = .789), highlighting the specificity of the relationship between awe and the cortical representation of ambivalence.

In AutoML-based multivariate analysis, adding 𝜙*_ambivalent_* metrics to a model predicting awe ratings with 14 behavioral variables improved the adjusted *R*^2^ value by 7.2% (see **Fig 4e**). This supports the significant contribution of ambivalence representation distinctiveness to the subjective experience of awe. Furthermore, 𝜙*_ambivalent_* demonstrated higher predictive power than other behavioral metrics in the predictive model (see **Fig 4f**), suggesting that the distinctiveness of cortical representations for ambivalent feelings may be a key factor driving individual differences in awe experiences.

These results indicate that individual differences in the distinctiveness of cortical representation of ambivalent feelings specifically account for the awe experience.

### The delta oscillation in the frontal areas mainly distinguishes different valence representation in the latent neural space

Lastly, we investigated which EEG features were important for contrasting different valence states when CEBRA constructed latent neural spaces. We utilized explainable AI technique specialized for time-series data, Dynamask (Crabbé & Van Der Schaar, 2021). It estimates a feature’s importance by observing how perturbing it at a specific time point alters the CEBRA model’s latent space embedding. Specifically, Dynamask learns a weight matrix by perturbing feature values with nearby values based on a perturbation weight, producing a pseudo-embedding with a high mean-squared error from the original CEBRA embedding (see **Fig 5a**; see ‘Methods’ for details). If a perturbation weight is close to 1, it indicates that the feature is crucial for the CEBRA model in learning the latent space embedding (i.e., contrasting the valence category at that time point from other valences). We independently learned the perturbation weight matrix for each participant’s clip data. By aligning the temporal axis of the weight matrix with participants’ valence dynamics ratings, we calculated the mean perturbation weight of each feature for every valence category. To calculate mean perturbation weights for each valence label, we used only data displaying significant silhouette coefficients for the corresponding valence label (e.g., in the case of ambivalent feelings, data colored in **Fig 4b**).

**Fig 5.**
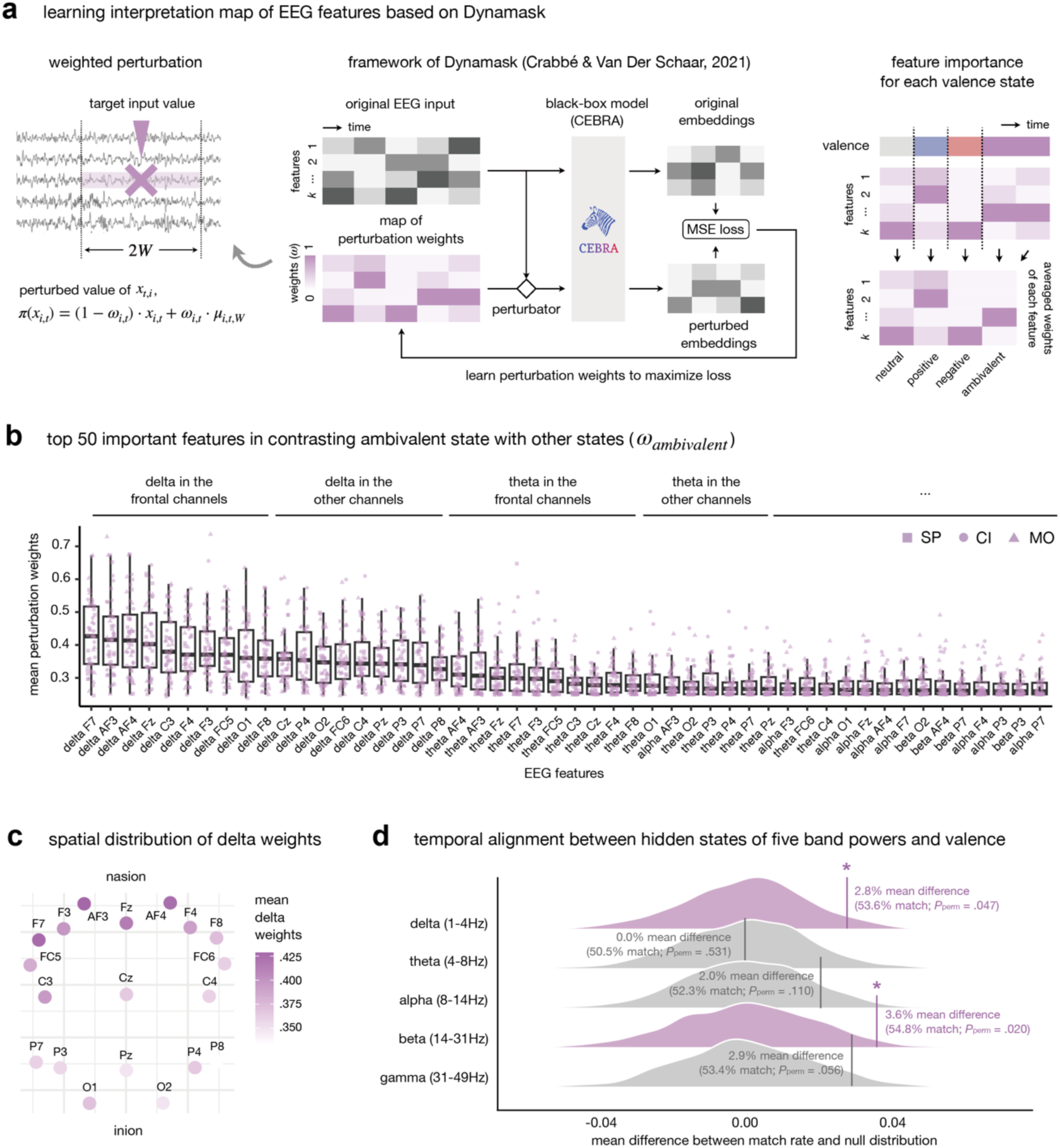
EEG features distinguishing different valence types. **a,** Schematic illustrating the use of Dynamask to learn an attribution map for EEG features. **b,** Top 50 most important features for differentiating ambivalent feelings from other valence categories. **c,** Spatial distribution of feature importance for delta power. **d,** Mean difference between the observed match rate and the null distribution for each frequency band, assessed through a permutation approach. The vertical line indicates the observed mean difference. Purple distributions represent statistically significant differences (*P*_perm_ < .05), while gray distributions are non-significant. Results are based on data from *N* = 27 participants included in the electrophysiological analysis.

Firstly, we confirmed that all Dynamask models reached plateau in learning performance for learning the weight matrix (see **Supplementary Fig 4**). Delta band power features had higher mean perturbation weights for ambivalent states than other frequency features (see **Fig 5b**), suggesting that delta oscillations may play a crucial role in differentiating ambivalent states from other valences in the brain. Within the delta band, frontal channels had relatively higher weights than channels in other areas (see **Fig 5c**), further implicating frontal delta oscillations in the processing and representation of ambivalent feelings. The same analysis for positive and negative states also showed that delta band power in the frontal channels consistently exhibited higher mean perturbation weights (see **Supplementary Fig 5**), indicating that frontal delta oscillations might be broadly involved in valence processing beyond ambivalent feelings.

To verify the importance of delta oscillation in valence representation, we conducted a post-hoc analysis using a hidden Markov model (HMM). We hypothesized that state changes in delta band power might temporally align with transitions in self-reported valence. We calculated the match rate between event boundaries in delta oscillations and key-pressed valence sequences and assessed its significance through a 1,000 times permutation test – calculating *P*_perm_ following the previous literature (Vaccaro et al., 2024). Neural boundaries from delta-related features significantly matched transitions in participants’ valence ratings above random chance (match rate = 53.6%; *P*_perm_ = .047), providing further support for a temporal link between delta oscillations and the valence dynamics. Among the other frequency band power features, only beta power showed a significant match rate (see **Fig 5d**). These findings suggest that delta oscillation in the frontal channels is crucial for distinguishing different valence states in the latent neural space.

## Discussion

Our study reveals that awe is fundamentally characterized by ambivalence, challenging traditional unidimensional models of valence. By combining immersive virtual reality, EEG, and advanced machine learning, we demonstrate that awe experiences are marked by ambivalent feelings at both behavioral and neural levels. Crucially, the ’cortical distinctiveness’ of ambivalence representation, captured through individualized latent neural spaces, significantly predicts the subjective intensity of awe. This novel metric quantifies how distinguishable ambivalent neural representations are from other valence states. Our findings establish a direct link between the neural processing of mixed emotions and the phenomenological experience of awe, offering new insights into how the brain integrates seemingly contradictory affective states. This approach not only illuminates the intricate nature of awe but also provides a novel framework for understanding and quantifying the neural basis of emotional complexity.

The significant association of the duration and intensity of ambivalent feelings with awe intensity further supports our hypothesis that awe is inherently an ambivalent experience. This finding aligns with previous cross-cultural research; for example, Western populations rated threat-awe-inducing images as having stronger ambivalence than stimuli evoking happiness or fear (Chaudhury et al., 2022). This consistency across cultures and stimuli suggests that the ambivalent nature of awe may be a fundamental feature of this emotion.

Our investigation of neural representations of ambivalence during awe revealed that aligned embeddings from individualized latent neural spaces successfully predicted valence dynamics across participants and stimuli, outperforming traditional methods like PCA and FAA. This superiority aligns with known limitations of FAA in valence indexing (Gable & Harmon-Jones, 2010; Harmon-Jones & Gable, 2018; Honk & Schutter, 2006; Wacker et al., 2003) and extends beyond PCA’s capabilities in detecting generalizable neural trajectories, such as motor control (Gallego et al., 2020; Safaie et al., 2023). Our deep neural net approach effectively captured idiosyncratic and shared valence representations, reflecting nonlinear brain-valence relationships (Aftanas et al., 1998; Berridge, 2019; Viinikainen et al., 2010) at the individual level. Notably, our method successfully disentangled affective neural representations from varied sensory inputs, demonstrating its robustness across diverse stimuli.

Our latent neural spaces revealed distinct cortical representations for ambivalent feelings, consistent with studies indicating that ambivalent feelings have unique neural patterns in cortical regions (Lettieri et al., 2019; Man et al., 2017; Vaccaro et al., 2020; Vaccaro et al., 2024). This finding seems to directly challenge the constructivists’ view that ambivalent feelings lack specific neural correlates (Barrett & Bliss-Moreau, 2009; Russell, 2017). Nevertheless, our findings of individual variability in these neural representations may integrate these conflicting arguments into a single framework. We observed individual variability in how neural representations of ambivalent feelings are cohesive in the latent space, which paradoxically supports the constructivist perspective that psychological factors (e.g., attentional focus or emotional granularity) influence how mixed emotions are experienced and processed (Hoemann et al., 2017). This dual finding - distinct neural patterns alongside individual variations - offers a novel framework for understanding complex emotions. Our deep neural network (CEBRA) approach, by capturing both aspects, potentially reconciles these divergent views in affective neuroscience, demonstrating the nuanced nature of ambivalence processing in the brain.

A key finding of our study is that the cortical distinctiveness of ambivalent states, quantified by our novel metric, strongly predicts self-reported awe intensity, outperforming traditional behavioral measures. Our approach, which considers the geometrical characteristics of neural representations, revealed relationships between ambivalent feelings and awe intensity that were overlooked by previous studies focused solely on activation levels in specific brain regions (Guan et al., 2019; Hu et al., 2017; Takano & Nomura, 2022). This underscores the value of our analytical method in uncovering the nuanced neural underpinnings of complex emotional experiences like awe.

The positive correlation between cortical distinctiveness of ambivalence and awe intensity aligns with the concept of “holistic meaning-making” in awe experiences (Bonner & Friedman, 2011; Dai et al., 2022; Ihm et al., 2019; Sawada et al., 2024). Awe is often characterized by encounters with stimuli that challenge our existing cognitive schemas, requiring us to accommodate new information and generate new understandings. This process may involve integrating initially evoked negative feelings, perhaps triggered by the vastness or novelty of the stimuli, with the positive feelings that emerge from successful comprehension and meaning-making. Likewise, awe is often described as a ‘self-transcendent’ experience, where two conflicting feelings – ‘self-diminishment’ and ‘connectedness’ are harmonized (Yaden et al., 2016; Yaden et al., 2019). Awe is thus based on integrating opposing feelings, creating an emotion beyond mere positivity and negativity. The distinctiveness of ambivalence-related neural representations observed in our study may reflect the consequence of this dynamic integration process, suggesting that the brain is actively processing both positive and negative aspects of the experience, ultimately contributing to a more personally meaningful and intense experience of awe.

Our analyses of Dynamask (Crabbé & Van Der Schaar, 2021) and HMM (Kumar et al., 2021) consistently highlighted the significance of frontal delta oscillations in distinguishing valence states. This aligns with previous neuroimaging findings on the role of frontal cortices such as the orbitofrontal cortex and anterior cingulate cortex in processing conflicting affective information (Levens & Phelps, 2010; Rolls & Grabenhorst, 2008; Simmons et al., 2006) and consistent activity patterns when individuals feel ambivalence during movie watching (Vaccaro et al., 2024). The prominence of frontal delta activity suggests its crucial role in integrating positive and negative feelings during awe.

These frequency band, encompassing both delta and low-theta ranges (below 5-6Hz), has been consistently linked to emotional processing and regulation in the literature. Frontal delta power increases during emotional memory retrieval (Brenner et al., 2014; Hutchison & Rathore, 2015; Nishida et al., 2009; Sopp et al., 2017), while the 1-3Hz range specifically decodes valence ratings to images(Shen et al., 2020). Theta burst stimulation, enhancing 5Hz band power in the dorsolateral prefrontal cortex, improves emotion recognition for ambiguous lexical and facial stimuli (Dumitru et al., 2020; Moulier et al., 2021), further supporting the role of low-frequency bands in emotional processing. Moreover, emotion regulation has been linked to 4Hz band power in frontal areas (Cavanagh & Shackman, 2015; Ertl et al., 2013); while individuals with impaired emotion regulation has been linked to decreased power in frequencies below 5Hz (Jiang et al., 2022). Indeed, in our HMM analysis, the synchrony of delta and beta features, which was aligned with real-time valence dynamics, has also been implicated in efficient emotion and stress regulation (Brooker et al., 2021; Myruski et al., 2022; Phelps et al., 2016; Putman et al., 2012). These findings collectively suggest frontal delta oscillations’ critical involvement in constructing coherent emotional responses to the complex, often conflicting feelings elicited by awe-inspiring experiences, supporting our observations on the neural dynamics of ambivalence during awe.

Our findings, coupled with existing literature on frontal delta power and emotion regulation, suggest a potential role for emotion regulation processes in mediating ambivalent feelings during awe experiences. Ambivalent feelings are often thought to stem from the reappraisal process (Vaccaro et al., 2020; Van Tilburg et al., 2018), where initial affective responses are re-evaluated through memory retrieval or knowledge associated with opposite valences. While we did not directly measure emotion regulation strategies in this study, previous studies have shown that reappraisal predicts awe ratings for recalled memories (Chirico et al., 2024; Chirico et al., 2021). Additionally, studies on affect dynamics show emotion regulation types correlate with the duration of negative feelings triggered by stimuli (Van Mechelen et al., 2013; Verduyn et al., 2009; Verduyn et al., 2011), but not the intensity, of negative feelings (Brans & Verduyn, 2014). These findings raise the intriguing possibility that the extended duration of ambivalent feelings we observed during awe experiences might originate from the engagement of implicit reappraisal processes. To further explore this hypothesis, future research could employ cognitive task analysis or think-aloud protocols during VR experiences. Such approaches would allow for direct assessment of reappraisal strategies during awe, illuminating how individual emotion regulation patterns influence ambivalent feelings and awe at both behavioral and neural levels.

Our study has limitations. We acknowledge that our sample comprised only Asian participants, potentially overestimating the role of ambivalent feelings in awe. Western individuals report fewer ambivalent feelings in awe experiences compared to Asians (Nakayama et al., 2020; Stellar et al., 2024). While previous research has found that Western individuals also report higher ambivalence for awe-inducing stimuli than single-valence stimuli (Chaudhury et al., 2022), future studies should investigate cross-cultural differences in the neural representation of ambivalence during awe. Additionally, using real-time valence ratings through keypresses might have inadvertently influenced participants’ emotional experiences (Larsen & Fredrickson, 1999). However, recent naturalistic affective science studies have validated and applied moment-by-moment emotion ratings during watching movies (Belfi et al., 2019; Lettieri et al., 2019; Vaccaro et al., 2024) or listening to music (McClay et al., 2023). Finally, our contrastive learning approach does not directly address how different valence types might share common neural substrates. Investigating these shared representations, particularly in brain regions associated with both pleasure and pain (Lee et al., 2024), could provide a more comprehensive understanding of the neural dynamics underlying ambivalence.

In conclusion, our study provides insights into the ambivalent nature of awe and its neural underpinnings. By demonstrating the link between ambivalent feelings, their distinct cortical representation, and the subjective intensity of awe, our findings advance the field of affective neuroscience and contribute to a more nuanced understanding of this complex and powerful emotion. This work lays the groundwork for further investigations into the cognitive and neural processes that underpin awe, paving the way for a more comprehensive understanding of its potential benefits for mental well-being.

## Methods

### Participants

We recruited 50 healthy young adult Koreans enrolled in psychology courses at Seoul National University for this study. Participants were excluded if they met any of the following criteria: (1) currently taking psychiatric medication, (2) history of psychiatric treatment, (3) left-handedness, (4) vestibular neuritis or balance disorder, (5) visual acuity before correction less than 0.2, (6) non-Korean native speakers, and (7) consumption of alcohol or use of hair rinse 24 hours before the experiment. Following these exclusion criteria, data from 43 participants were completely collected in the analyses (23 females; *M*_age_ = 20.2 years, *SD*_age_ = 1.7 years). Seven participants were excluded due to technical issues (*N* = 3), discontinuance due to motion sickness (*N* = 2), and lack of fidelity in online valence ratings (i.e., showing accuracy below 75% in the training session for valence keypress; *N* = 2). The sample size was chosen to be comparable to recent naturalistic affective neuroscience studies – e.g., (Lettieri et al., 2019; Meer et al., 2020; Vaccaro et al., 2024). Participants provided written informed consent before the experiment, and all procedures were approved by the Institutional Review Board of Seoul National University.

### Experimental paradigm

#### VR clip design

We collaborated with a professional filmmaker to design four audio-integrated 360° immersive videos using Unreal Engine (version 5.03). Each video lasted 120 seconds. Three of the videos were designed to evoke awe: *Space* (Meer et al.), *City* (CI), and *Mountain* (MO), while the other one, *Park* (PA), served as a control stimulus, designed not to include any awe-related factors. To investigate whether ambivalence is consistently observed in various awe experiences, we varied (1) the clip themes, (2) the VR cues of awe-related two dimensions – perceived vastness and the need for accommodation, and (3) the perceptual features across the awe-inducing videos.

Firstly, considering that awe is most intensively triggered by massive landscapes (Chirico et al., 2018; Keltner & Haidt, 2003; Shiota et al., 2007; Yaden et al., 2019), we differentiated the semantic theme of the scenery: SP featured supernatural landscapes (i.e., black holes and planets in space), CI showcased urban landscapes (i.e., cityscape viewed from the top of skyscrapers), and MO depicted natural one (i.e., mountain scenery).

Secondly, following the qualitative framework of the ref(Chirico et al., 2018)to design VR videos that effectively elicit awe, we aimed to represent the two key dimensions of awe, perceived vastness and a need for accommodation, through different cues in each video. Each video was designed so that perceivers would first experience vastness during the initial 60 seconds and then feel a need for accommodation in the latter 60 seconds. For example, in SP, participants watched a giant black hole approaching, consuming everything and ultimately drawing them in, followed by the sudden appearance of Earth from space. Different sub-components were applied to realize each dimension across clips. Perceived vastness can be induced through perceptual (e.g., ‘width’ and ‘height’) and conceptual cues (e.g., ‘complexity’)(Chirico et al., 2017; Chirico et al., 2018). MO was designed to evoke vastness through perceptual width, CI through height, and SP through conceptual complexity. For the need for accommodation, we introduced surprise cues in each video around the 60-second mark as a trigger of accommodation (Chirico et al., 2017; Chirico et al., 2018), tailored to the context of each video to ensure that the cause of surprise did not overlap across videos. The design of three awe-inducing clips is summarized in **Supplementary Fig 1a**.

Lastly, to prevent awe from being driven by specific perceptual factors, we intentionally composed the three awe videos with different audiovisual information. We synchronized visual content with ambient sounds using open-source audio samples from Freesound (https://freesound.org) and GarageBand (version 10.4.6). To verify our design, we calculated three perceptual features known to predict perceivers’ emotional responses – color hue (Dael et al., 2016; Suk & Irtel, 2010), brightness (Kurt et al., 2017), and loudness (Thao et al., 2019) – every second for each stimulus and visualized their time-course dynamics. We qualitatively confirmed that each video exhibited very different temporal dynamics for all features (see **Supplementary Fig 1b**).

To validate the awe elicitation, we conducted a preliminary study with 28 independent young adult Koreans (five females; *M*_age_ = 20.2 years, *SD*_age_ = 1.9 years), who rated awe intensity using the Awe Experience Scale (AWES)(Yaden et al., 2019) after watching each clip in VR. Participants reported significantly higher awe ratings for three awe clips than the control clip, with large effect sizes (SP vs. PA: Cohen’s *d* = 2.466, *P*_FDR_ = 8’10^-13^; CI vs. PA: Cohen’s *d* = 2.193, *P*_FDR_ = 6’10^-12^; MO vs. PA; Cohen’s *d* = 1.52, *P*_FDR_ = 1’10^-6^).

#### Baseline self-report

Before the experiment, participants provided information on their sex, age, baseline mood states, and dispositional traits of feeling awe. Baseline mood states were assessed using the Korean version of the Positive and Negative Affect Schedule (PANAS) validated by (Lim et al., 2010). No participants showed exceptional mood states beyond 1.5 ’ interquartile range (*M*_positive_ = 33.581, *SD*_positive_ = 6.284; *M*_negative_ = 23.140; *SD*_negative_ = 6.331). Dispositional awe was measured using the awe-related items from the Dispositional Positive Emotions Scale (DPES)(Shiota et al., 2006) translated into Korean (*M* = 29.628, *SD* = 6.626). The Korean-translated DPES items showed acceptable reliability (Cronbach’s a = .827) but mixed results in validity (model fit of the original factor structure: CFI = .946, RMSEA = .127).

#### Procedures

Participants sat on a sofa in a noise-isolated room and wore an Enobio 20 EEG device (Neuroelectrics) and a Quest 2 VR headset (Oculus). After checking EEG signal quality, the experiment proceeded as follows: baseline EEG recording, keypress training, VR watching task, post-trial measurement, and a break. Firstly, participants’ resting EEG signals were recorded for 120 seconds with their eyes closed (baseline recording). These resting signals were used to normalize signals recorded during the movie-watching trials. Secondly, they practiced real-time valence keypress reporting (keypress training). Participants were explicitly asked to report their valence feelings in real-time keypress with the following auditory instruction: *“While watching the video, please report your affective state by pressing a number pad: ‘1’ for positive, ‘2’ for ambivalent - feeling positive and negative feelings at the same time, and ‘3’ for negative feelings. If you do not feel any affective feelings, please do not press anything. If a specific feeling persists, continue to press and hold the corresponding key. It is important to report your subjective reactions rather than which emotions the video intends to elicit”*. Participants practiced this for 60 seconds using sentences describing affective responses to prevent confusion about which key they should press. Then, participants watched four VR clips in a pseudorandom order, reporting their valence using online keypress (VR watching task). After each trial, they reported awe intensity, overall valence, arousal, and motion sickness using controllers (post-trial measurement). Awe intensity was measured by the Korean-translated AWES (Yaden et al., 2019), valence by Evaluative Space Grid (Larsen et al., 2009), arousal by a conventional 9-point Likert scale (Bradley & Lang, 1994), and motion sickness by a single 7-point Likert scale item. The Korean-translated AWES demonstrated acceptable psychometric properties (Cronbach’s a = .928; model fit of the original factor structure: CFI = .881, RMSEA = .079).

Participants took a 30-second break with their eyes closed after each trial (break).

### EEG data acquisition

#### EEG recording and preprocessing

We recorded EEG signals using 19 dry electrodes: AF3, AF4, F7, F3, Fz, F4, F8, FC5, FC6, C3, Cz, C4, P7, P3, Pz, P4, P8, O1 and O2 with Neuroelectrics Enobio 20 based on the international 10-20 system. Ground and reference electrodes were attached to the right earlobe. The embedded software in the Enobio system assessed signal quality using three levels: good, medium, and bad. We ensured that no electrodes displayed bad signals before starting the signal acquisition. We adopted the automated preprocessing pipeline validated by (Delorme, 2023). EEG signals for each trial were time-locked to the initiation of the video, excluding the last three seconds to avoid end-of-task effects (e.g., loss of attention or emotional confounding). High-pass filtering above 0.5 Hz and Artifact Subspace Reconstruction were performed. Unlike the original pipeline, we used interpolation to maintain consistent recording lengths across participants and trials rather than excluding time windows with poor signal quality. We conducted independent component analysis-based artifact rejection to remove noise components, such as eye or head movements, with over 90% probability. The preprocessed signal for each trial was normalized by subtracting the average resting signal value for each channel. All preprocessing was performed using the “EEGLAB” plugin (Delorme & Makeig, 2004) in MATLAB (version 2021a).

#### Short time/Fast Fourier transform

With preprocessed and normalized EEG signals, we performed short time Fourier transform (STFT) and fast Fourier transform (FFT) to calculate the spectral power of five frequency bands for each channel: delta (1-4Hz), theta (4-8Hz), alpha (8-14Hz), beta (14-31Hz), and gamma (31-49Hz). For STFT, a Hanning window with a 500-sample window size (i.e., 1 sec) and a 250-sample hop size was applied. Participants’ valence keypresses were embedded as event markers in EEG signals, categorizing EEG samples into one of four valence categories. The valence label of each 500-sample window after STFT was defined as the mode of the corresponding samples’ valence labels. FFT was also performed to calculate overall spectral powers marginalized across the whole time series. Using the Welch method, we calculated the power spectral density for each EEG channel and then integrated it over the specific frequency range described above to determine the band power. For relative band power, we normalized the power within each band by the total power across all frequencies. The “scipy” package (Virtanen et al., 2020) in Python (version 3.8) was used for STFT and FFT.

### Behavioral analysis

#### Univariate linear mixed-effects model analysis

Using data from 43 participants, we first assessed the univariate association between AWES ratings and 14 behavioral features measured before, during, and after each trial: sex, age, PANAS positive score, PANAS negative score, DPES awe score (before trial), duration of positive, ambivalent, negative, and neutral feelings (during trial), arousal, motion sickness score, and intensity of positive, ambivalent, and negative feelings (after trial). The duration of each valence type was calculated as the ratio of keypresses for that valence type to the total running time of each clip. Intensity was calculated based on the Evaluative Space Grid responses: positivity (x-axis value), negativity (y-axis value), and ambivalence (minimum value between positivity and negativity, following previous literature (Berrios et al., 2015; Chaudhury et al., 2022; Ersner-Hershfield et al., 2008). Firstly, we performed two-sided paired t-tests to examine statistical differences in AWES scores, duration, and intensity of each valence type, and arousal between the three awe clips and the control clip at *P*_FDR_ < .05. Next, to evaluate the explanatory power of the 14 metrics, we fit linear mixed effect models with each regressor and two random intercepts for participants and clips using the “lmerTest” package (Kuznetsova et al., 2015). Assumptions of normality were examined using the “DHARMa” package (Hartig, 2018). We confirmed that the distribution of residuals did not significantly deviate from a normal distribution using the Kolmogorov-Smirnov test (all *P*s > .05). All variables were standardized before the statistical analysis. All statistical analyses were conducted in R studio (version 2023.03.1+446).

#### Multivariate machine learning analysis

Next, we conducted machine learning-based predictive modeling with 14 behavioral variables for AWES scores as multivariate analysis, considering potential non-linear interactions among features. Using “h2o” package (LeDell & Poirier, 2020), we split the dataset into training and test sets with a 4:1 ratio and conducted 5-fold cross-validation. A total of 22 models were constructed, and we selected the best model based on the lowest fold-averaged test RMSE value. In the model selection, models that did not provide feature importance information (e.g., stacked ensemble models) were excluded for interpretability. As a result, the gradient boost model (GBM) was chosen as the best model. Its predictive performance was compared to a baseline ridge linear regression model without any interaction terms for metrics: RMSE, MAE, MSE, and *R*^2^. To identify the most influential features, we calculated feature importance and shapley values for each variable. All machine learning analyses were performed in R studio (version 2023.03.1+446).

### Electrophysiological analysis

#### Construction of individualized latent neural space for each clip

Among 43 participants in the behavioral analysis, we excluded 16 individuals due to poor-quality EEG signals even after preprocessing (*N* = 6) and lack of ambivalent keypresses in at least one trial (i.e., who reported ambivalent feelings for less than 5% of the total duration across all videos; *N* = 10). The quality of preprocessed signals was visually inspected. The primary objective of the electrophysiological analysis was to investigate whether ambivalent feelings during awe have distinct cortical representations and to evaluate their relationship with the intensity of awe. Given that our behavioral analysis identified the duration of ambivalent feelings as the most salient predictor of awe ratings, such exclusion is inevitable and expected not to introduce sampling bias regarding the measurement of awe ratings.

Using 27 participants’ EEG signals and valence keypress in the three awe clip trials, we constructed a latent neural space for each individual and clip (i.e., a total of 81 latent spaces) using the “CEBRA” package (Schneider et al., 2023). CEBRA employs supervised contrastive learning to extract latent embeddings from the input data (i.e., STFT-processed EEG signals here), maximizing the attraction of EEG samples with the same valence labels and repelling those with different labels. As the dimensionality of latent spaces was elusive, we fitted CEBRA models for each participant-clip pair across dimensions ranging from one to nine. The following hyperparameters were applied: batch_size = length of STFT EEG signals, model_architecture = ‘offset-10 model’, number_of_hidden_units = 38, learning_rate = .001, the number_of_iterations = 500, and hybrid = False.

#### Validation of latent neural spaces with predictive tasks

To test whether the individualized latent spaces hold significant information generalizable across different individuals and clips, we conducted a pairwise prediction task using 2 tasks ’ 3 conditions design. For the ‘across participants’ task, a classifier trained on Participant A’s latent neural space embeddings and valence labels was used to predict the valence keypress of Participant B’s embeddings for the same clip. For the ‘across clip’ task, a classifier trained on clip X’s embeddings and labeled valence types was used to predict the valence of clip Y’s embeddings within the same participant. Predictions were evaluated under three conditions: (1) a baseline null test with shuffled training valence keypress labels (‘random’), (2) prediction using personalized latent neural embeddings without any alignment between train and test embeddings (‘not-aligned’), and (3) prediction using aligned embeddings between train and test embeddings (‘aligned’). Neural alignment motivates the exploration of commonality among individual latent neural spaces (Gallego et al., 2020; Safaie et al., 2023). Here, canonical correlation analysis (CCA) between train and test embeddings was employed using the “sklearn” package (Pedregosa et al., 2011).

In these prediction tasks, six participants who pressed all four valence labels for at least 5% of the total duration across three clips were selected to facilitate the multi-label classification. A k-nearest neighbors (kNN) classifier with a neighborhood parameter of 15 was used (i.e., the nearest odd number to the square root of the input embedding length following the conventional heuristic). Considering the imbalance in valence keypress labels, prediction performance was evaluated using the weighted F1 scores. A two-sided paired t-test was conducted to compare predictive performances across the three conditions *P*_FDR_ < .05. Construction of latent neural spaces and predictive analyses were performed in Python (version 3.8).

#### Dimensionality selection for the latent neural spaces

We selected the optimal dimensionality of latent neural spaces based on two assumptions: (1) The space shares the same dimensionality across individuals and clips. (2) The space displays as good as predictive performances across individuals and clips with other neural spaces based on higher dimensionality, even with fewer dimensions. The second assumption was based on the idea that generalizability, as reflected in predictive performance, should be the criterion for determining the canonical dimensions in dimensionality reduction techniques (Cunningham & Yu, 2014). However, it also acknowledges that such predictive performance may be overestimated as the number of dimensions increases (Cunningham & Yu, 2014; Diaconis & Freedman, 1984). We calculated each dimensionality’s mean weighted F1 score n predictive tasks and performed hierarchical clustering analysis to group dimensions with similar prediction performance. The number of clusters was chosen for the highest silhouette coefficients, and the lowest dimension in the highest-performing cluster was selected. Dimensions 6, 7, 8, and 9 formed the high-performance cluster for the across-participant task, and dimensions 7, 8, and 9 for the across-clip task. Thus, the 7D space condition was chosen as the canonical dimensionality. Hierarchical clustering analysis was conducted using the “cluster” package (Maechler et al., 2013) in R studio (version 2023.03.1+446).

#### Comparing CEBRA-, PCA-, and FAA-driven latent space embeddings

For a fair comparison of the predictive power of our CEBRA-based latent neural embeddings, we compared its performance with PCA and FAA-driven embeddings. First, to extract PCA embeddings, we input the STFT-processed EEG features and computed latent embeddings with dimensions ranging from one to nine. We identified the 6-dimensional embedding as the optimal latent space due to its modest predictive performance with the fewest dimensions. Second, FAA embeddings were defined as the difference in alpha band power between the F4 and F3 channels for each timepoint in the STFT-featured EEG sequence. These two channels were selected based on previous studies (Brzezicka et al., 2017; Quaedflieg et al., 2016; Van Der Vinne et al., 2017). The prediction tests with the 2 × 3 design were conducted using these three types of embeddings. Test performances under the three conditions – random, not aligned, and aligned – were compared across CEBRA, PCA, and FAA-based embeddings. Additionally, within the CEBRA embeddings, performances were compared across the different conditions. To test the statistical differences in weighted F1 scores, we conducted a two-sided paired t-test at *P*_FDR_ < .05. PCA was performed using the “sklearn” package (Pedregosa et al., 2011) in Python (version 3.8).

#### Assessing significance of cortical valence representation

Using the chosen CEBRA-based 7D latent space embeddings, we measured the segregation of ambivalence-related EEG samples from other valence samples using silhouette coefficients. While InfoNCE loss value could quantify the contrast performance of CEBRA as well, it lacks scaling and valence type-specific calculations, so we used silhouette coefficients instead. We computed the average silhouette coefficient for ambivalence-labeled EEG samples for each participant-clip latent space. Since silhouette coefficients can be overestimated for clusters in the latent space constructed in a supervised manner, we assessed its statistical significance through a permutation test. We randomly shuffled valence keypresses and trained the CEBRA model with the original STFT-featured EEG signals and permuted valence sequence based on the identical hyperparameter set. Average silhouette coefficients of ambivalence-labeled EEG samples were calculated from the trained pseudo-embeddings, and we obtained its null distribution by repeating this 1,000 times. The *P*-value calculated from the permutation test, *P*_perm_, is formulated as follows:

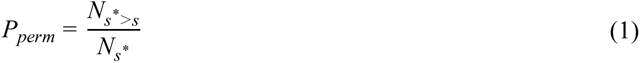

where *s\** and *s* are average silhouette coefficients calculated from permuted and original latent embeddings, and *N_s*_* is the number of *s\** in the null distribution. All *P*_perm_ values were FDR corrected. We performed the same analysis with positive and negative valence clusters in the latent neural space.

#### Quantifying ‘cortical distinctiveness’ of each valence in the latent neural space

We developed a metric called ‘cortical distinctiveness’, 𝜙*_k_,* indicating how distinguishable the reference cluster *k* is from the other valence clusters in the latent valence-cortical space. 𝜙*_k_* is defined as:

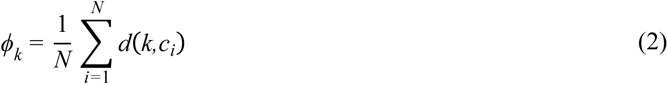

where *N* is the number of other clusters, *c_i_* is the *i*-th valence cluster, and *d*(*k, c_i_*) is the cosine distance between the cluster *k* and *c_i_*. We applied cosine distance as a cluster distance metric instead of other conventional metrics (e.g., Euclidean distance), considering that latent CEBRA embeddings are distributed on the hypersphere space. We initially measured *d*(*k, c_i_*) based on the average cluster distance. The average distance between the cluster *k* and *c_i_* is calculated as the mean cosine distance between each point in *k* to every point in *c_i_* as follows:

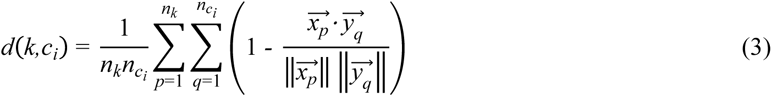

where *n* is the sample size of the corresponding cluster, and 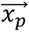 and 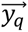 are the vector samples in each cluster. As a sensitivity check for cluster distance metric, we also calculated 𝜙*_k_* based on the medoid cluster distance. Medoid distance measures the distance between clusters by calculating the cosine distance between ‘medoid samples’ of each cluster, which shows the closest average cosine distance with samples within each cluster. To test the predictive power of cortical distinctiveness of each valence type for AWES scores, we applied the same univariate and multivariate analysis framework as described in the “Behavioral analysis” section.

#### Inference of feature importance in CEBRA using Dynamask

Due to the black-box nature of CEBRA, it was challenging to directly evaluate which STFT-processed EEG features were crucial for building latent neural spaces in CEBRA (i.e., contrasting valence types in each time point). To address this issue, we applied “Dynamask” (Crabbé & Van Der Schaar, 2021), a perturbation-based explainable AI techniques, to infer attribution maps from the trained CEBRA models. Dynamask learns perturbation weights, *w*, for each feature at every time point to generate pseudo-embeddings with maximal MSE value compared to the original embeddings with the slightest perturbation. Here, the *i*-th input feature at the timepoint *t*, *x_i,t_* is perturbed to p(*x_i,t_*) as a weighted sum of its own value and the average value in the time window it belongs to, formulated by the following equation:

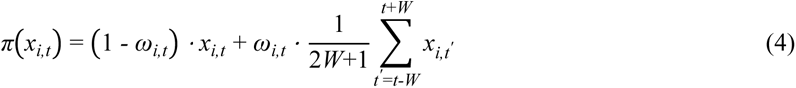

where *W* is the time window size. *w_i,t_* = 1 indicates a significant alteration of the CEBRA embeddings upon replacement, implying importance in contrasting valence labels at the given time point. With a receptive field of 10 samples, *W* = 5 was set to align the working behavior of Dynamask with CEBRA’s.

We obtained *w* matrices for each EEG feature from data showing significant silhouette coefficients for ambivalence clusters (*P*_perm_ < .05) using the following parameters: keep_ratio = 0.1, n_epoch = 2,500, initial_mask_coef = 0.5, size_reg_factor_init = 0.5, size_reg_factor_dilation = 100, time_reg_factor = 0, learning_rate = 0.03, and momentum = 0.9. By aligning the time point of STFT-processed EEG features and valence keypress sequence, we calculated *w*_ambivalent_ of each feature by averaging *w* values of each feature for time points labeled as ambivalent states in the valence keypress. Consequently, *w*_ambivalent_ provides an indirect measure of feature importance to contrast ambivalence-labeled EEG samples from other valence-labeled EEG samples to construct latent valence-cortical space. The same analyses were conducted for positive and negative valence states with data displaying significant silhouette coefficient for each state, respectively.

#### Post-hoc analysis of feature importance with hidden Markov model

We conducted a post-hoc analysis using the hidden Markov Model (HMM) to confirm Dynamask-driven attribution weights of each feature in distinguishing different valence states within the latent neural space. Particularly, Dynamask revealed that the power of the delta band held greater importance than the power of other frequency bands. Based on this, we hypothesized that hidden states of delta features would be temporally aligned with individual valence dynamics and tested this hypothesis using an HMM. For all HMM analyses, we used the “brainiak” package (Kumar et al., 2021).

We divided the STFT-processed EEG features into frequency bands to generate five input groups – delta, theta, alpha, beta, and gamma – for each participant and clip. These inputs were independently fitted to an HMM, estimating the time points of neural boundaries corresponding to the number of valence transitions reported via keypress. Thus, the event number, a hyperparameter of the HMM, was informed by the participants’ reported valence keypress for the clip. Boundaries within ± 3 seconds of the actual valence transition time points were considered a ‘match’, and the ‘match rate’ was calculated as the ratio of matched boundaries to the total number of boundaries.

Following the framework of previous literature (Vaccaro et al., 2024), we assessed the statistical significance of the match rates for the five frequency bands across all participants and clips. This framework’s advantage is that it accounts for variability in the number of valence transitions reported by each participant for each clip. The approach involves the following steps: First, for each participant and frequency band input, maintain the estimated intervals of neural boundaries but shuffle them randomly, comparing these to the actual valence transition time points to compute a pseudo-match rate. This process is repeated 1,000 times to generate a null distribution of match rates. Second, calculate the difference between the actual match rate and the mean of the null distribution for each frequency band group for each participant-clip data. Averaging these differences across participants and clips yields the average mean difference for feature group *k*, denoted as *M_k_*. Third, repeat this process 1,000 times using the null distribution of each participant-clip data to derive a null distribution of 1,000 mean differences between permuted match rates and the mean of the null distribution. Denote the *i*-th permuted mean difference for feature group *k* as 𝑄*_k_^(i)^*. Last, calculate the *P* value, *P*_perm_, for *M_k_* using the following equation:

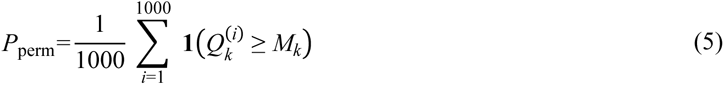

## Acknowledgements

This work was supported by the National Research Foundation of Korea (NRF) grant funded by the Korea government (MSIT) (No. 2021R1C1C1006503, RS-2023-00266787, RS-2023-00265406, RS-2024-00421268), by Creative-Pioneering Researchers Program through Seoul National University (No. 200-20240057), by Semi-Supervised Learning Research Grant by SAMSUNG (No.A0426-20220118), by Identify the network of brain preparation steps for concentration Research Grant by LooxidLabs (No.339-20230001), by Institute of Information & communications Technology Planning & Evaluation (IITP) grant funded by the Korea government (MSIT) [NO.RS-2021-II211343, Artificial Intelligence Graduate School Program (Seoul National University)] by the MSIT (Ministry of Science, ICT), Korea, under the Global Research Support Program in the Digital Field program (RS-2024-00421268) supervised by the IITP (Institute for Information & Communications Technology Planning & Evaluation), by the National Supercomputing Center with supercomputing resources including technical support (KSC-2023-CRE-0568) and by the Ministry of Education of the Republic of Korea and the National Research Foundation of Korea (NRF-2021S1A3A2A02090597), and by Artificial intelligence industrial convergence cluster development project funded by the Ministry of Science and ICT (MSIT, Korea) & Gwangju Metropolitan City.

## Author contributions

J.Y. and J.C. conceived the study and designed the behavioral experiment. J.Y. and S.Y.O. designed the VR movie stimuli and experimental protocol. J.Y. and D.D.H. developed the code. J.Y. performed behavioral and EEG data analysis. J.Y., D.D.H., and J.C. interpreted the results and drafted the manuscript.

---

**Supplementary Fig 1.**
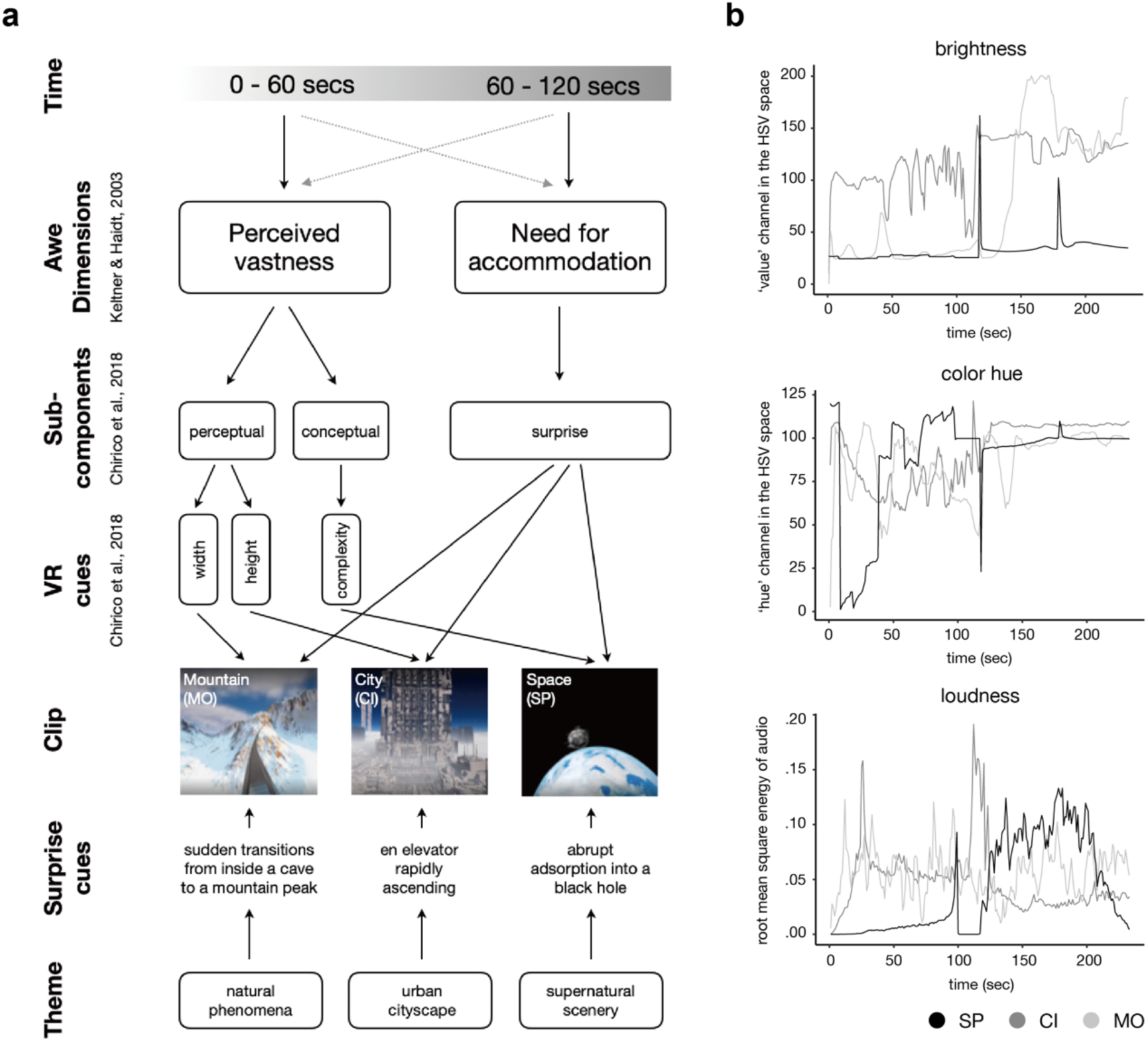
Design of VR clips to induce diverse awe experience using distinct awe-related cues and sensory profiles. **a,** We designed the three awe-inducing VR clips to evoke distinct awe experiences while incorporating key dimensions of awe, each implemented through different sub-components in the VR environment. Additionally, each video was intentionally crafted with a unique. The common structure (i.e., ‘awe dimensions’) was based on the core dimensions of awe, which was formulated by Keltner and Haidt (2003). To ensure that each awe dimension was distinctly represented in the VR environment, we applied the qualitative framework of Chirico et al. (2018), which suggests distinct sub-components of perceived vastness and relevant VR cues. **b,** The heterogeneous temporal patterns of visual and acoustic features in three awe-inducing clips.

**Supplementary Fig 2.**
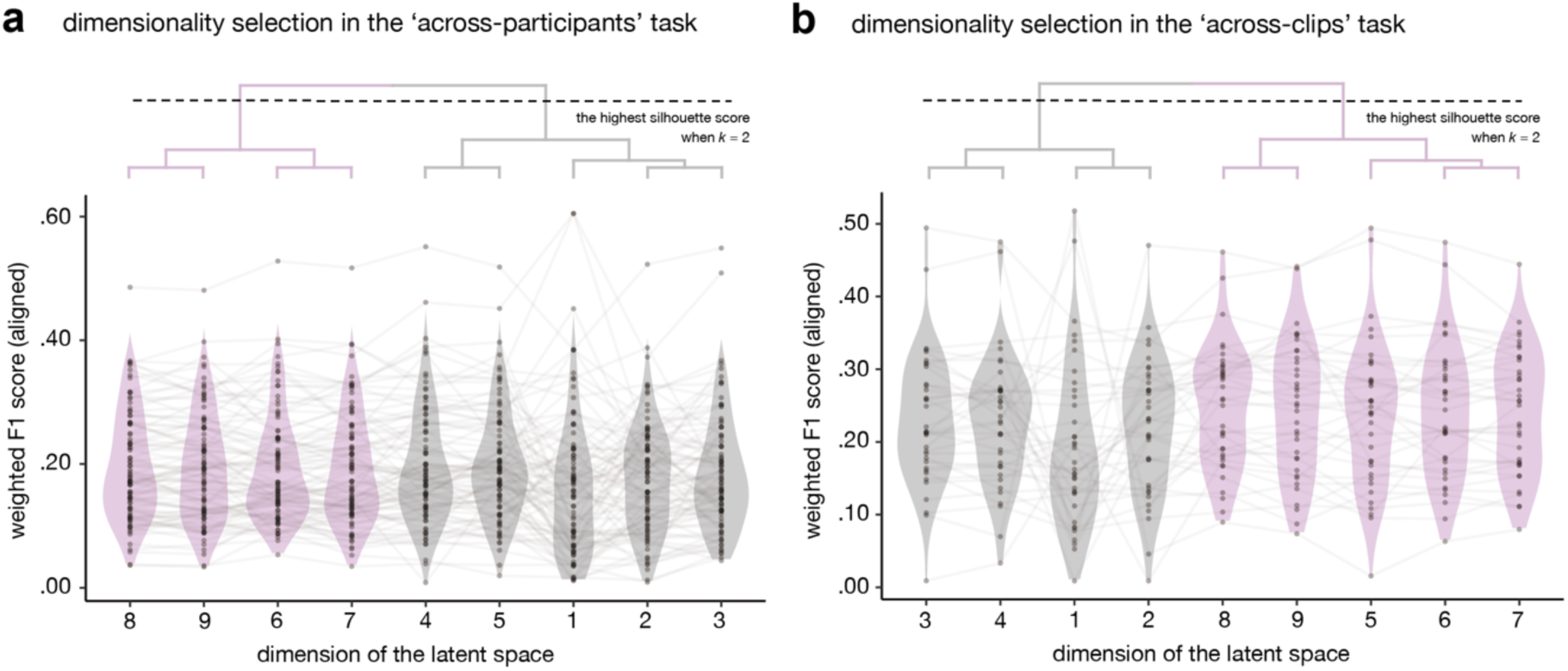
Dimensionality selection for PCA-driven latent neural space. **a,** The test performances in the ‘across-participants’ task. Hierarchical clustering analysis identified clusters with high performance in decoding other participants’ valence dynamics (purple violin plots). **b,** The test performances in the ‘across-clips’ task.

**Supplementary Fig 3.**
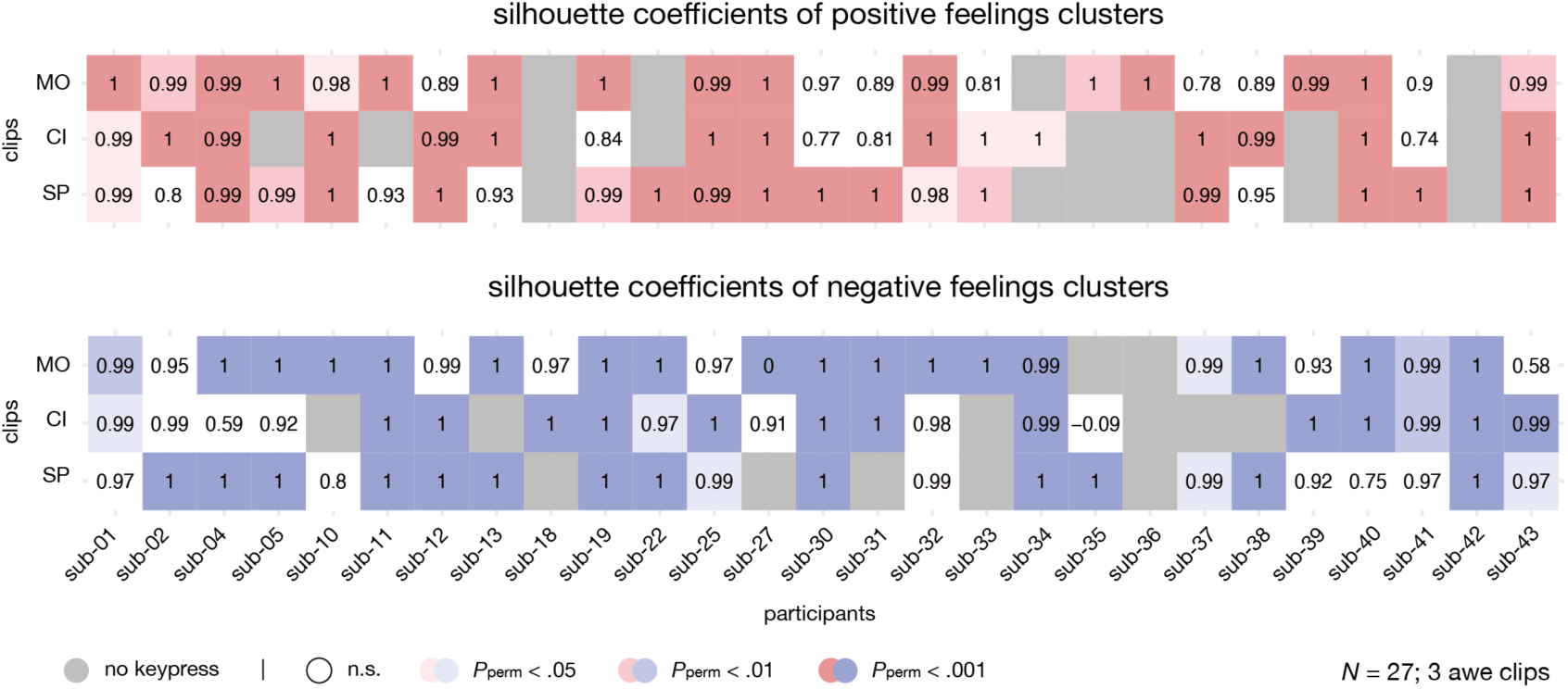
Silhouette coefficients of positive and negative valence clusters. Silhouette coefficients of positive (red) and negative (blue) feelings clusters within individualized latent cortical spaces. Color intensity indicates the statistical significance of the silhouette coefficients. Some participants have a gray cell (i.e., not respond positive or negative feelings in the keypress) because 27 participants included in this analysis were chosen in terms of whether they responded ambivalent feelings for more than 5% of the total duration across all awe-inducing clips, rather than positive or negative feelings.

**Supplementary Fig 4.**
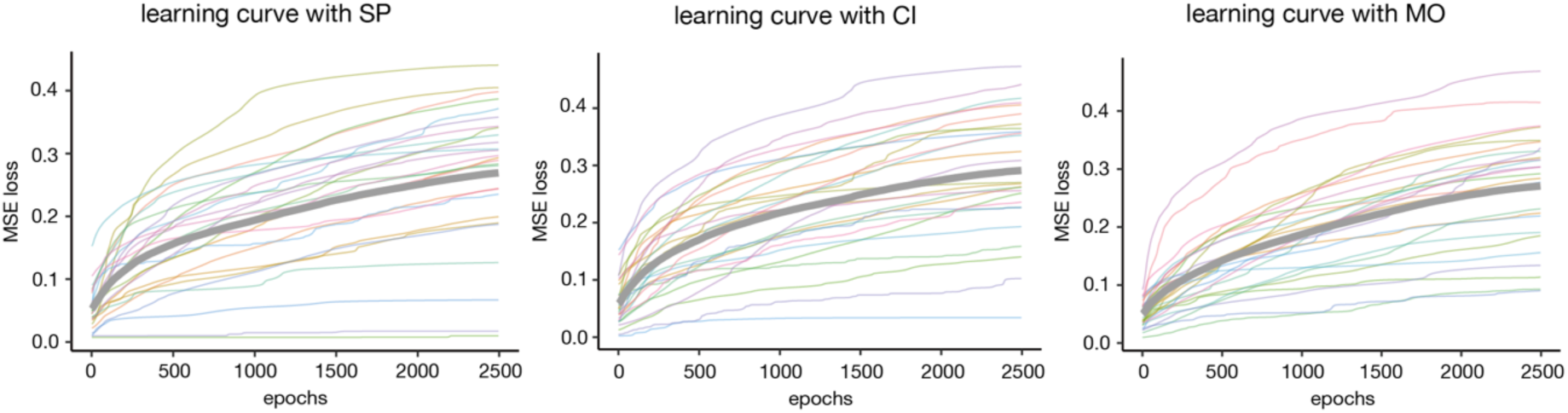
Learning curves of Dynamask to learn attribution map. Dynamask learns perturbation weights of EEG features for every time point in terms of the importance in constructing latent cortical space through CEBRA (i.e., distinguishing the valence at the given timepoint with other ones). Thus, it aims to extract perturbation weights matrix so that resultant pseudo-embeddings from perturbed inputs have large mean-squared error (MSE) with the original latent space embeddings as much as possible. Each colored curve denotes its learning curve of respective participants’ data. Thick gray curves visualize the average loss values at every epoch.

**Supplementary Fig 5.**
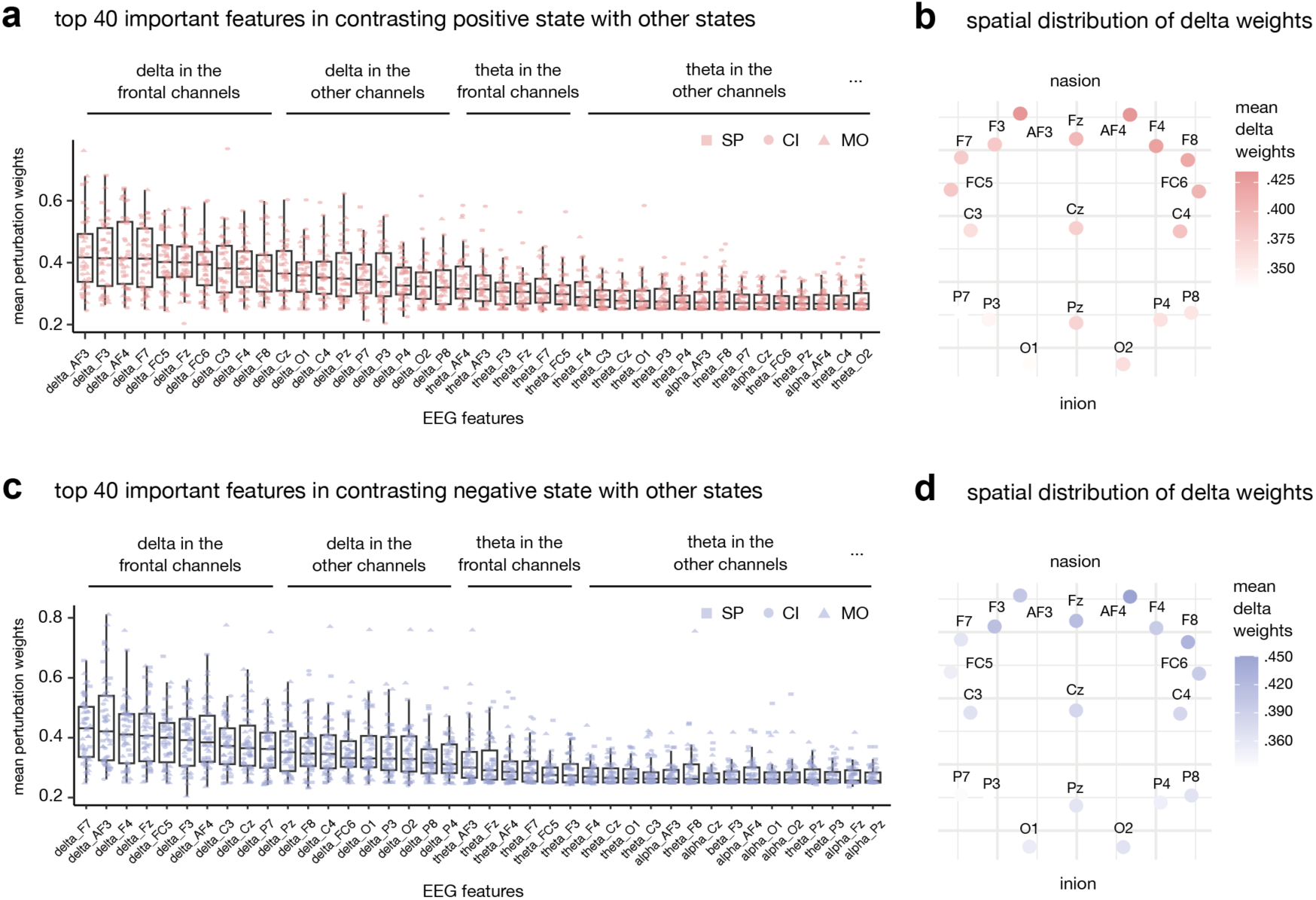
Perturbation weights of EEG features for positive and negative feelings. **a,** Top 40 most important features for differentiating positive feelings from other valence categories. **b,** Spatial distribution of mean perturbation weights of delta-related features for positive feelings. **c,** Top 40 most important features for differentiating negative feelings from other valence categories. **b,** Spatial distribution of mean perturbation weights of delta-related features for negative feelings.

**Supplementary Table 1.**
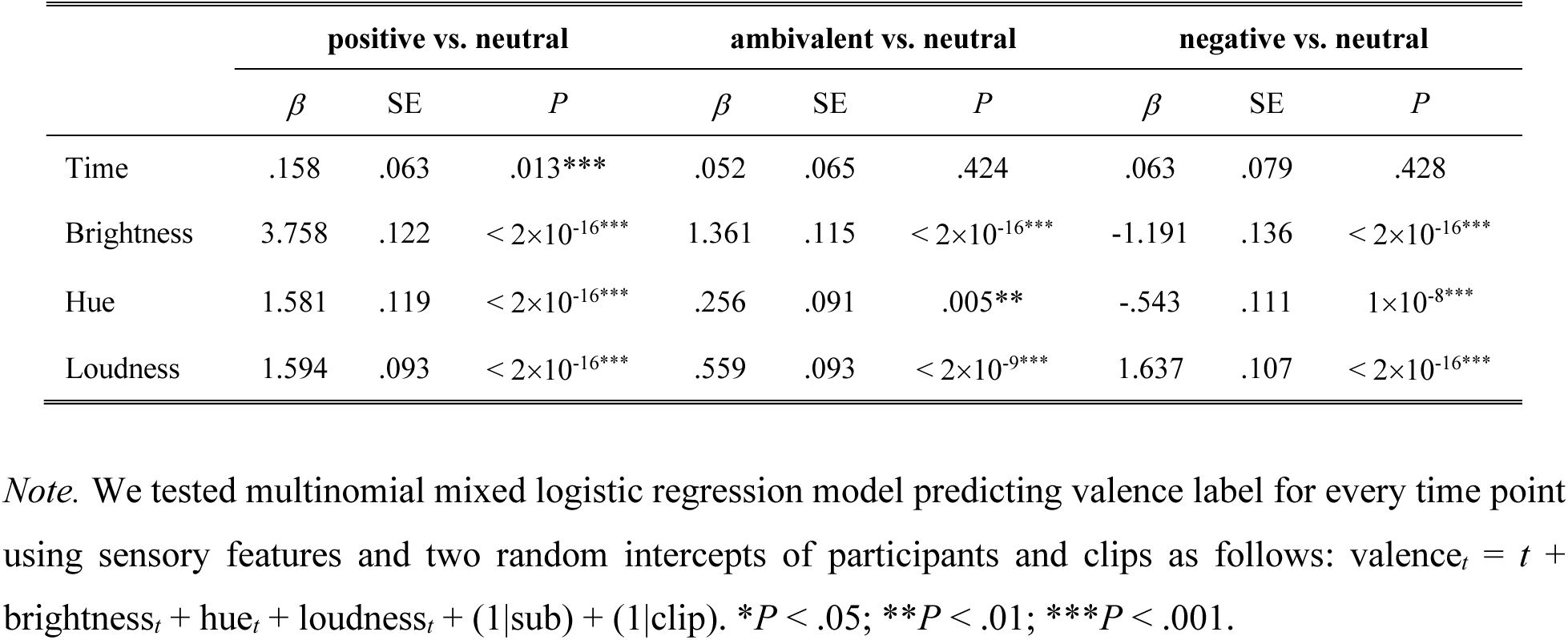
Associations between affect-related sensory features of clips and participants’ valence dynamics.

|  | positive vs. neutral |  |  | ambivalent vs. neutral |  |  | negative vs. neutral |  |  |
| --- | --- | --- | --- | --- | --- | --- | --- | --- | --- |
| | $\beta$ | SE | $P$ | $\beta$ | SE | $P$ | $\beta$ | SE | $P$ |
| Time | .158 | .063 | .013*** | .052 | .065 | .424 | .063 | .079 | .428 |
| Brightness | 3.758 | .122 | $< 2 \times 10^{-16}$*** | 1.361 | .115 | $< 2 \times 10^{-16}$*** | -1.191 | .136 | $< 2 \times 10^{-16}$*** |
| Hue | 1.581 | .119 | $< 2 \times 10^{-16}$*** | .256 | .091 | .005** | -.543 | .111 | $1 \times 10^{-8}$*** |
| Loudness | 1.594 | .093 | $< 2 \times 10^{-16}$*** | .559 | .093 | $< 2 \times 10^{-9}$*** | 1.637 | .107 | $< 2 \times 10^{-16}$*** |

*Note.* We tested multinomial mixed logistic regression model predicting valence label for every time point using sensory features and two random intercepts of participants and clips as follows: $\text{valence}_t = t + \text{brightness}_t + \text{hue}_t + \text{loudness}_t + (1|\text{sub}) + (1|\text{clip})$ . \* $P < .05$ ; \*\* $P < .01$ ; \*\*\* $P < .001$ .

**Supplementary Table 2.** Statistical differences in valence and arousal ratings across clips.

|  | SP vs. PA |  |  |  | CI vs. PA |  |  |  | MO vs. PA |  |  |  |
| --- | --- | --- | --- | --- | --- | --- | --- | --- | --- | --- | --- | --- |
|  | *t* | *df* | *d* | *P*<sub>FDR</sub> | *t* | *df* | *d* | *P*<sub>FDR</sub> | *t* | *df* | *d* | *P*<sub>FDR</sub> |
| **Valence** |  |  |  |  |  |  |  |  |  |  |  |  |
| ambivalent (int) | 2.957 | 42 | .552 | .005** | 2.946 | 42 | .576 | .005** | 5.118 | 42 | .961 | 2×10<sup>-5***</sup> |
| ambivalent (dur) | 3.685 | 42 | .775 | .001*** | 3.372 | 42 | .770 | .002** | 5.179 | 42 | 1.218 | 2×10<sup>-5***</sup> |
| positive (int) | 1.427 | 42 | .266 | .352 | -.443 | 42 | .092 | .660 | 1.206 | 42 | .225 | .352 |
| positive (dur) | -3.316 | 42 | .667 | .003** | -3.618 | 42 | .795 | .002** | -3.178 | 42 | .634 | .003** |
| negative (int) | -.295 | 42 | .059 | .769 | 2.562 | 42 | .431 | .021* | 3.024 | 42 | .536 | .013* |
| negative (dur) | .421 | 42 | .082 | .676 | 4.329 | 42 | .839 | 1×10<sup>-4***</sup> | 5.243 | 42 | 1.085 | 1×10<sup>-5***</sup> |
| **Arousal** | 5.810 | 42 | 1.151 | 7×10<sup>-7***</sup> | 7.404 | 42 | 1.433 | 6×10<sup>-9***</sup> | 9.301 | 42 | 1.777 | 3×10<sup>-11***</sup> |

*Note.* ‘int’ and ‘dur’ denote intensity and duration of the corresponding feelings, respectively. \**P*<sub>FDR</sub> < .05;
\*\**P*<sub>FDR</sub> < .01; \*\*\**P*<sub>FDR</sub> < .001.

**Supplementary Table 3.** Statistical differences in key demographics and awe ratings between the participants included and excluded in the electrophysiological analysis.

|  | Included<br>(*N* = 27) | Excluded<br>(*N* = 16) | *P*<sub>FDR</sub> |
| --- | --- | --- | --- |
| **Demographics** |  |  |  |
| male; *N* (%) | 13 (48.1) | 7 (43.8) | .733 |
| age years; *M*(*SD*) | 20.2 (1.7) | 20.3 (1.7) | .918 |
| **AWES ratings** |  |  |  |
| for SP; *M*(*SD*) | 4.7 (0.6) | 4.4 (0.7) | .559 |
| for CI; *M*(*SD*) | 4.6 (0.5) | 3.8 (1.0) | .070 |
| for MO; *M*(*SD*) | 4.3 (0.8) | 4.1 (1.0) | .733 |

*Note.* Statistical difference of the categorical variable (i.e., male) between groups was assessed by Brunner-Munzel test. For the other variables, we applied two-sided t-test. All *P* values were FDR corrected.

**Supplementary Table 4.** Statistical differences in predictive performances of latent neural space embeddings across conditions and analytic approaches.

|  | across participants |  |  |  | across clips |  |  |  |
| --- | --- | --- | --- | --- | --- | --- | --- | --- |
| | *t* | *df* | *d* | $P_{\text{FDR}}$ | *t* | *df* | *d* | $P_{\text{FDR}}$ |
| **within CEBRA embeddings** |  |  |  |  |  |  |  |  |
| *aligned vs. random* | 10.641 | 89 | 1.122 | $5 \times 10^{-17}^{***}$ | 3.005 | 35 | .501 | .015* |
| *aligned vs. not-aligned* | 5.200 | 89 | .548 | $2 \times 10^{-6}^{***}$ | 1.699 | 35 | .283 | .147 |
| *not aligned vs. random* | 2.366 | 89 | .249 | .020* | .620 | 35 | .103 | .540 |
| **between embeddings** |  |  |  |  |  |  |  |  |
| *aligned* |  |  |  |  |  |  |  |  |
| CEBRA vs. PCA | 10.371 | 89 | 1.093 | $2 \times 10^{-16}^{***}$ | 3.320 | 35 | .553 | .006** |
| CEBRA vs. FAA | 10.576 | 89 | 1.115 | $6 \times 10^{-17}^{***}$ | 4.840 | 35 | .807 | $8 \times 10^{-5}^{***}$ |
| *not-aligned* |  |  |  |  |  |  |  |  |
| CEBRA vs. PCA | 1.830 | 89 | .193 | .106 | .360 | 35 | .060 | .721 |
| CEBRA vs. FAA | 1.898 | 89 | .200 | .091 | .452 | 35 | .075 | .782 |
| *random* |  |  |  |  |  |  |  |  |
| CEBRA vs. PCA | -.379 | 89 | .040 | .706 | -.410 | 35 | .068 | .721 |
| CEBRA vs. FAA | .269 | 89 | .028 | .789 | -.278 | 35 | .046 | .782 |

Note. \* $P_{\text{FDR}} < .05$ ; \*\* $P_{\text{FDR}} < .01$ ; \*\*\* $P_{\text{FDR}} < .001$ .

## Notes

### Competing Interest Statement

The authors have declared no competing interest.

### Summary of Updates

Abstract and Discussion parts were revised to enhance readability.

